# Sex-specific and Developmental Effects of Early Life Adversity on Stress Reactivity are Rescued by Postnatal Knockdown of 5-HT_1A_ Autoreceptors

**DOI:** 10.1101/2024.01.22.576344

**Authors:** Rushell Dixon, Lauren Malave, Rory Thompson, Serena Wu, Yifei Li, Noah Sadik, Christoph Anacker

**Affiliations:** Division of Systems Neuroscience, Department of Psychiatry, Columbia University, and Research Foundation for Mental Hygiene, Inc. (RFMH), New York State Psychiatric Institute (NYSPI), New York, NY, 10032, USA; Columbia University Institute for Developmental Sciences, Research Foundation for Mental Hygiene, Inc. (RFMH)/New York State Psychiatric Institute (NYSPI), Department of Psychiatry, Columbia University Irving Medical Center (CUIMC), New York, NY, 10032, USA; Columbia University Stem Cell Initiative (CSCI), Columbia University Irving Medical Center (CUIMC), New York, NY, 10032, USA

**Keywords:** Hypothalamus-Pituitary-Adrenal Axis, Monoamines, Glucocorticoids, Neural Circuits, Childhood Trauma

## Abstract

Early Life Adversity (ELA) predisposes to stress hypersensitivity in adulthood, but neurobiological mechanisms that protect from the enduring effects of ELA are poorly understood. Serotonin 1A (5HT_1A_) autoreceptors in the raphé nuclei regulate adult stress vulnerability, but whether 5HT_1A_ could be targeted to prevent ELA effects on susceptibility to future stressors is unknown. Here, we exposed mice with postnatal knockdown of 5HT_1A_ autoreceptors to the limited bedding and nesting model of ELA from postnatal day (P)3-10 and tested behavioral, neuroendocrine, neurogenic, and neuroinflammatory responses to an acute swim stress in male and female mice in adolescence (P35) and in adulthood (P56). In females, ELA decreased raphé 5HT neuron activity in adulthood and increased passive coping with the acute swim stress, corticosterone levels, neuronal activity, and corticotropin-releasing factor (CRF) levels in the paraventricular nucleus (PVN) of the hypothalamus. ELA also reduced neurogenesis in the ventral dentate gyrus (vDG) of the hippocampus, an important mediator of individual differences in stress susceptibility, and increased microglia activation in the PVN and vDG. These effects of ELA were specific to females and manifested predominantly in adulthood, but not earlier on in adolescence. Postnatal knockdown of 5HT_1A_ autoreceptors prevented these effects of ELA on 5HT neuron activity, stress reactivity, neurogenesis, and neuroinflammation in adult female mice. Our findings demonstrate that ELA induces long-lasting and sex-specific impairments in the serotonin system, stress reactivity, and vDG function, and identify 5HT_1A_ autoreceptors as potential targets to prevent these enduring effects of ELA.

## INTRODUCTION

Early life adversity (ELA) is a major risk factor for cognitive and emotional disturbances, and ∼30% of mental illnesses in adulthood can be attributed at least in part to ELA.^1^ Despite this high prevalence and the potent and long-lasting effects of ELA on mental health, we still lack a comprehensive understanding of the biological mechanisms that determine individual differences in vulnerability and resilience to adverse early experiences.^2^

Enhanced vulnerability for psychopathology following ELA has been associated with altered physiological and emotional reactivity to stressors that are experienced later in life.^3–5^ Accordingly, dysregulated hypothalamus-pituitary-adrenal (HPA) axis responses have consistently been found in individuals with a history of ELA.^5,6,7,8^ Rodent models recapitulate ELA effects on HPA axis dysregulation and have shown that maternal separation or early life limited bedding and nesting (LBN) conditions lead to higher corticosterone (CORT) levels at baseline and following acute stress in adulthood.^9–16^ These ELA effects on HPA axis hyperactivity have been associated with heightened activity of the paraventricular nucleus (PVN) of the hypothalamus, which leads to increased synthesis and release of corticotropin-releasing factor (CRF), in turn triggering the release of adrenocorticotropic hormone (ACTH) and thereby stimulating the release of CORT from the adrenals.^17–19^

We and others have previously shown that chronically elevated CORT levels reduce adult hippocampal neurogenesis in the dentate gyrus (DG)^20–22^ - a region of the hippocampus that regulates behavioral and neuroendocrine responses to stress.^23,24^ Accordingly, ELA causes alterations in the structure and cellular composition of the rodent DG, including reduced dendritic arborization, neurogenesis, granule cell numbers, and DG volume.^25–31^ Abnormalities in hippocampal structure and volume are also found in human subjects at high familial risk for depression^32,33^ and in children with a history of childhood adversity.^31,34–36^ Hippocampal impairments resulting from ELA, including reduced neurogenesis, may thus predispose to heightened stress responsivity later in life.

Both hypercortisolemia and inflammation are frequently observed in subsets of patients with depression.^37–40^ Even though CORT is known for its immunosuppressive effects, chronically elevated CORT levels are associated with increased inflammation, a phenomenon that has been attributed to glucocorticoid resistance and reduced sensitivity of the glucocorticoid receptor (GR) following its chronic activation by CORT.^41^ Pro-inflammatory effects of ELA on the brain are supported by studies showing higher numbers of activated microglia in the mouse and human hippocampus,^42,43,44^ as well as disrupted microglia function in the developing PVN, which promote HPA axis hyperactivity following acute stress in adulthood.^17^ ELA-dependent alterations in microglia activation may thus contribute to heightened stress reactivity, and increased inflammation has indeed been suggested to mediate an association between ELA and neurobiological correlates of mental illness.^45,46^

The extent to which ELA determines vulnerability to future stressors, HPA axis reactivity, and psychopathology can be influenced profoundly by genetic predisposition and biological sex.^47–52^ The serotonin 1A receptor (5HT_1A_) has been implicated in the regulation of neuroendocrine responses to stress in juvenile and adult animals,^53,8–12^ and the C(-1019)G polymorphism in the promoter region of the human *htr1a* gene is associated with depression and antidepressant treatment resistance.^13,14^ Previous studies have developed conditional transgenic knockout mice to probe the functional role of 5HT_1A_ receptors in specific brain regions.^54–56^ These studies have shown that germline deficient 5HT_1A_ knockout causes ‘anxiety-like’ behavior, an effect that can be reversed by rescuing 5HT_1A_ heteroreceptor expression in the hippocampus and cortex.^54^ 5HT_1A_ autoreceptors in the raphé nuclei of the brainstem inhibit the activity of 5HT neurons, thereby reducing 5HT release at terminal projection sites in limbic and cortical projection areas, including the hippocampus and PVN.^56^ Indeed, brain-wide deficiencies in 5HT increase stress vulnerability in rodents,^57^ while 5HT_1A_ autoreceptor knockdown confers stress resilience.^56^ Whether 5HT_1A_ autoreceptors also mediate differences in vulnerability to ELA effects is not known.

Previous work has comprehensively characterized behavioral impairments following ELA in adolescence and adulthood.^50,51,58^ However, neurobiological and molecular mechanisms underlying vulnerability and resilience to ELA effects across development remain largely unexplored. In this study, we tested whether 5HT_1A_ autoreceptors determine individual differences in ELA effects on stress reactivity, and how these effects develop throughout adolescence and early adulthood in male and female mice. We focused our neurobiological investigation on the hippocampus and PVN, two key regulators of the HPA axis in which 5HT release during stress is associated with avoidance behavior and active stress coping.^59^ We used the Forced Swim Test (FST) as a means to induce an acute stress response in adulthood and find that ELA-exposed females show reduced 5HT neuron activity, increased passive stress coping behavior, elevated CORT levels, PVN activity and CRF levels, reduced adult hippocampal neurogenesis, and increased neuroinflammation in the PVN and ventral DG. These impairments are specific to female mice and develop in adulthood, but not earlier on in adolescence. Postnatal knockdown of 5HT_1A_ autoreceptors prevented these effects of ELA on 5HT neurons, stress reactivity, neurogenesis, and neuroinflammation, indicating that restoring 5HT signaling through 5HT_1A_ autoreceptor knockdown can protect from the effects of ELA on stress hypersensitivity. Our findings shed new light on the biological mechanisms underlying sex-specific vulnerability to ELA and provide novel leads into potential future targets for intervention.

## MATERIALS AND METHODS

### Experimental mice

Procedures were conducted in accordance with the US National Institutes of Health (NIH) Guide for the Care and Use of Laboratory Animals and the New York State Psychiatric Institute (NYSPI) Institutional Animal Care and Use Committee (IACUC). Mice were housed 3–5 per cage with *ad libitum* access to food and water on a 12:12h light/dark cycle. To induce postnatal knockdown of 5HT_1A_ autoreceptors in raphé 5-HT neurons, we used Pet1-tTS;tetO-1A mice containing the tetracycline operator (tetO) in the promoter region of the *htr1a* gene and the tetracycline-dependent transcriptional suppressor (tTS) under control of the 540Z Pet-1 promoter fragment (Pet1-tTS), which is specific to raphé serotonin neurons.^56^ In the presence of doxycycline (DOX), raphé 5HT_1A_ levels are indistinguishable between mice heterozygote for the *Pet1-tTS* transgene (Pet1-tTS^+^) and mice without the *Pet1-tTS* transgene (Pet1-tTS^-^). Removal of DOX causes suppression of *htr1a* expression in heterozygote Pet1-tTS^+^ mice (from here on referred to as “5HT1A K.D.”) but not in Pet1-tTS^-^ mice (from here on referred to as “WT”).^56^ Nulliparous Pet1- tTS^-^ females were bred with heterozygote Pet1-tTS^+^ males and maintained on DOX diet. DOX was removed three days before parturition. Female and male Pet1-tTS^-^ (WT) and Pet1-tTS^+^ pups (5HT1A K.D.) were raised on regular chow. Please see *Supplemental Information* for a comprehensive description of experimental set up.

### Limited Bedding and Nesting (LBN)

Litters were exposed to standard-rearing conditions or to the limited bedding and nesting (LBN) model, which causes fragmented and unpredictable maternal care as a form of ELA.^15^ Briefly, dams and their newborn litters were placed on a wire mesh grid and provided with 1/3 of nesting material and 1/3 of bedding material from P3-10.^15,60^ Mice were returned to standard-rearing conditions on P10. Standard-reared control cages were changed at P3 and P10 but otherwise left undisturbed. All litters were weaned at P21.

### Behavioral Testing

Please see *Supplemental Information* for a complete description of behavioral procedures.

### Corticosterone (CORT) EIA assay

Peripheral blood was obtained by submandibular blood draw at baseline 3 days before the Forced Swim Test (FST), and again 30 min post-FST. Blood was centrifuged at 3000rpm for 15 min and serum extracted and stored at -80°C. CORT was quantified using EIA assays (ArborAssays, Ann Arbor, MI).^61^

### RNAscope

Please see *Supplemental Information* for a complete description of RNAscope procedures.

### Immunohistochemistry

Brains were collected 1h following FST to assess cFos, Dcx, CD68, and Iba1. Mice were anesthetized with ketamine/xylazine and transcardially perfused with saline and ice-cold 4% paraformaldehyde (PFA). Brains were post-fixed overnight in 4% PFA at 4 °C, cryoprotected in 30% sucrose for 48 hours, frozen in optimum cutting temperature (OCT) compound, and sectioned at 50 μm along the dorsal-ventral axis of the hippocampus.^61^ See also *Supplemental Information*.

### Fluorescence Microscopy & Cell Quantification

Please see *Supplemental Information* for a complete description.

### CRF quantification by Western Blot

PVN tissue micropunches were collected from fresh-frozen brain tissue and protein extracted using Neuronal Protein Extraction Reagent (N-PER^TM^, Thermo Scientific) for determination of CRF protein levels by Western Blot. See also *Supplemental Information*.

### Statistics

Detailed statistical results for all experiments are reported in **Supp. Table 1**. Data was confirmed to be normally distributed using the Shapiro–Wilk test. Co-variate analyses were conducted using SPSS. All other analyses were conducted using GraphPad Prism 9. Behavior data were analyzed using Three-Way repeated measure (RM) Analysis of Variance (ANOVA) to assess ELA x genotype x time interactions in the FST, and Two-Way ANOVA to assess ELA x genotype interactions of average immobility in the FST and time in the open arms in the EPM. Three-Way ANOVA was used to assess ELA x genotype x post-FST CORT levels and body weights. RNAscope, Immunohistochemistry, and Western Blot data was analyzed using Two-Way ANOVA to test ELA x genotype interactions. Tukey *post hoc* tests were used in all experiments where applicable.

## RESULTS

### ELA and 5HT_1A_ autoreceptor knockdown exert opposing effects on 5HT neuron activity

To determine if ELA effects on stress reactivity can be rescued by reducing 5HT_1A_ autoreceptor expression, we used a transgenic mouse model to suppress 5HT_1A_ autoreceptors on raphé 5HT neurons throughout postnatal development (Pet1-tTS;tetO-1A mice) (**Fig. 1a**).^56,62^ To characterize the time course of 5HT_1A_ knockdown, we exposed litters to the limited bedding and nesting (LBN) model of ELA from P3 to P10 and collected brains for RNAscope analysis of 5HT_1A_ autoreceptor expression in TPH2^+^ raphé neurons at P3, P10, P35, and P56 (**Fig. 1b**). We find that 5HT_1A_ K.D. reduces the number of 5HT_1A_-positive TPH2-expressing 5-HT neurons in the raphé of females and males starting at P3 and lasting until P56 (**Fig. 1c-e**). ELA or sex did not significantly affect the extent of this knockdown (**Fig. 1d,e; Supp Fig. 1**).

**Figure 1:**
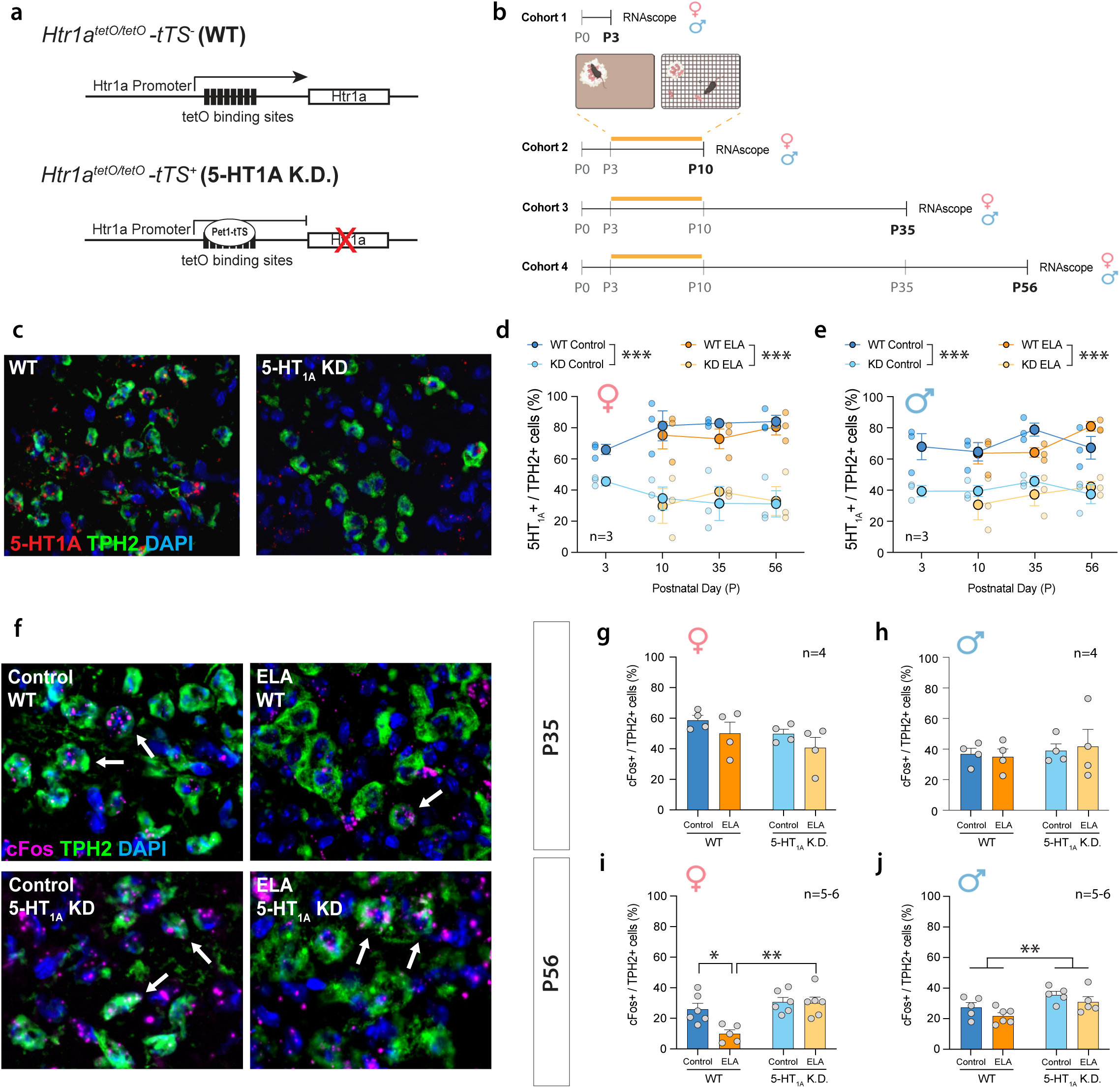
Opposing effects of ELA and 5HT1A autoreceptor knockdown on adult 5HT neuron activity. **a.** Schematic of the genetic knockdown of 5-HT1A autoreceptors in Pet1-tTS^-^ mice (“WT”) vs Pet1-Tts^+^ (“5HT1A K.D.”). **b.** Experimental timeline of ELA (P3-10) and tissue collection for RNAscope. **c.** Representative images of RNAscope for 5HT1A and TPH2 in WT and 5HT1A K.D. mice. **d.** Characterization of 5HT1A K.D. in females at P3, 10, 35, and 56. **e.** Characterization of 5HT1A K.D. in males at P3, 10, 35, and 56. **f.** Representative images of RNAscope for cFos and TPH2 in P56 females. Arrows indicate overlap between cFos and TPH2. **g.** At P35, ELA or 5HT1A K.D. do not affect the number of cFos+/TPH2+ cells in females, or **h.** in males. **i.** At P56, ELA reduces the number of cFos+/TPH2+ cells in WT females while 5HT1A K.D. rescues this effect of ELA. Brackets indicate significant Tukey *posthoc* tests. **j.** 5HT1A K.D. increases the number of cFos+/TPH2+ cells in adult males. Bracket indicates significant main effect of 5HT1A K.D. Mean±SEM; *P<0.05, **P<0.01, ***P<0.001.

Because 5HT_1A_ autoreceptors regulate 5HT neuron activity, we next determined 5HT neuron activity in ELA-exposed mice with and without 5HT_1A_ knockdown using RNAscope for the immediate early gene, cFos, as a proxy marker for neuronal activity in TPH2^+^ raphé 5HT neurons. In adolescence (P35), neither ELA nor 5HT_1A_ K.D. affected the number of cFos^+^/TPH2^+^ neurons in female or male mice compared to standard-reared WT controls (**Fig. 1g,h**). In adulthood (P56), the number of cFos^+^/TPH2^+^ neurons was reduced in ELA-exposed females, indicating decreased 5HT neuron activity (**Fig. 1f,i**). ELA did not affect the number of cFos^+^/TPH2^+^ neurons in males (**Fig. 1j**). 5HT_1A_ K.D. rescued ELA effects on cFos^+^/TPH2^+^ neurons in females and increased the overall number cFos^+^/TPH2^+^ neurons in males (**Fig. 1i,j**).

Together, these findings indicate that ELA in the form of LBN reduces 5HT neuron activity in adult females and that this effect can be rescued by 5HT_1A_ K.D.

### Stress responses in adulthood are elevated by ELA and rescued by 5HT_1A_ autoreceptor knockdown

To test ELA effects on stress responses later in life, we raised mice under LBN or standard-rearing conditions and exposed separate cohorts of adolescent (P35) and adult mice (P56) to a forced swim test (FST) to initiate an acute stress response. We then quantified immobility time in the FST, which is commonly interpreted as an indicator for passive stress coping or despair-like behavior in mice. We also measured serum CORT levels to evaluate endocrine responses at baseline and following the acute swim stress.

In adolescence (P35, **Fig. 2a**), neither ELA nor 5HT_1A_ K.D. affected immobility in the FST (**Fig. 2b,c**) or CORT levels at baseline or after the acute FST stress in females (**Fig. 2d**). However, 5HT_1A_ K.D. reduced overall CORT levels compared to WT females (**Fig. 2d**; *see* **Supp. Table 1** for detailed statistical comparisons). ELA did also not affect immobility or CORT levels in adolescent males (**Fig. 2e-g**), but 5HT_1A_ K.D. reduced immobility in males regardless of rearing conditions (**Fig. 2e,f**).

**Figure 2:**
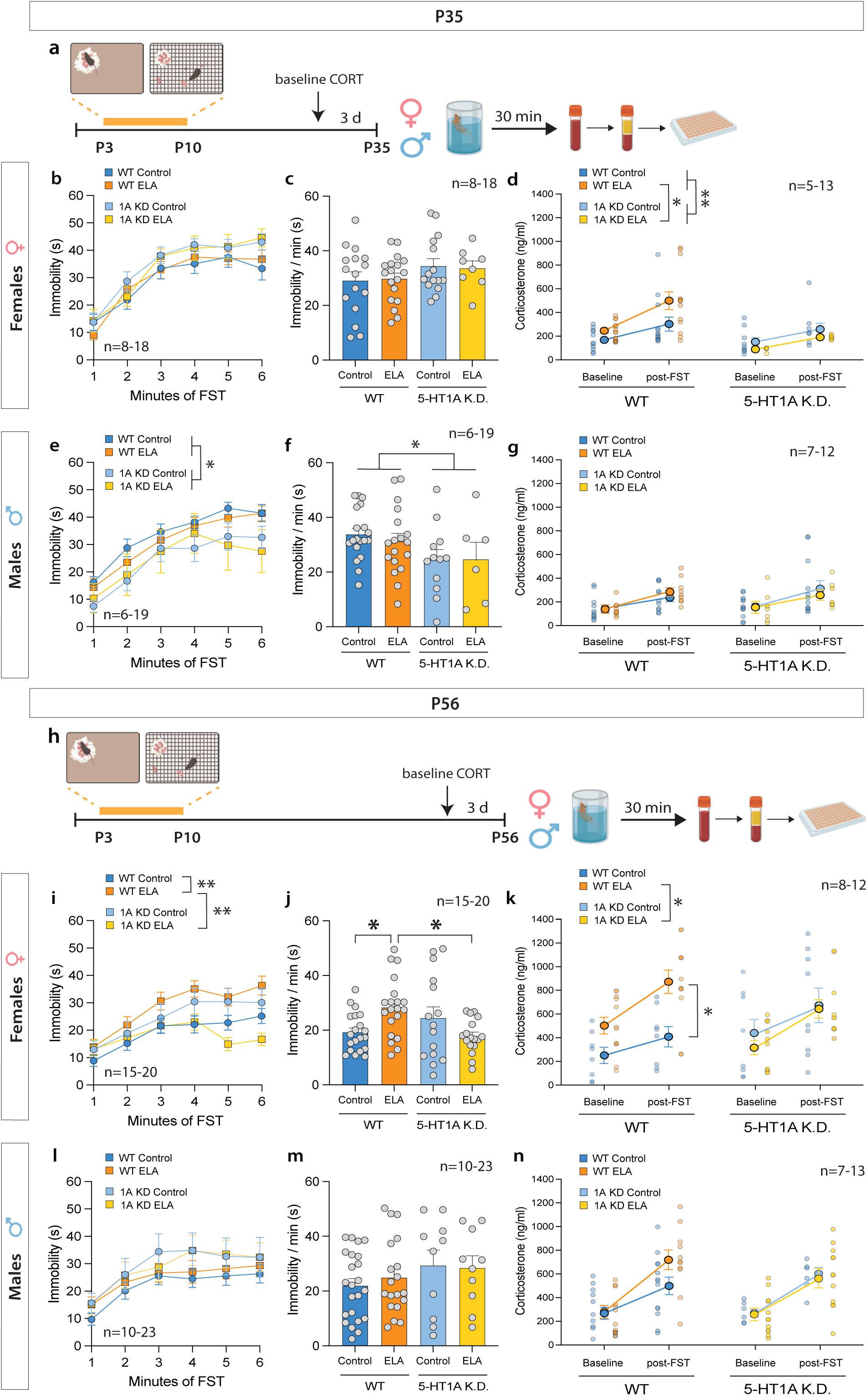
5-HT1A autoreceptor knockdown rescues ELA effects on behavioral and endocrine response to adult stress in females. **a.** Experimental timeline of ELA (P3-10), acute swim stress exposure, and blood collection in adolescence (P35). **b,c.** No difference in immobility time between adolescent WT and 5HT1A K.D. control vs. ELA-exposed females. **d.** 5HT1A K.D. reduces CORT levels compared to WT mice in adolescent females regardless of rearing condition. **e,f.** 5HT1A K.D. decreases immobility time in adolescent males. **g.** ELA or 5HT1A K.D. does not affect baseline or post-FST CORT responses in adolescent males. **h.** Experimental timeline of ELA (P3-10), acute swim stress exposure, and blood collection in adulthood (P56). **i,j.** ELA increases immobility in the FST in adult females compared to standard-reared control females. 5HT1A K.D. reduces immobility in ELA-exposed females. **k.** In WT females, ELA increases post-FST CORT responses compared to controls. No effect of ELA on CORT is observed in 5-HT1A K.D. females. **l,m.** No difference in immobility between adult WT and 5HT1A K.D. control vs. ELA-exposed males. **n.** Neither ELA nor 5-HT1A K.D. affect baseline or post-FST CORT responses in adult males. FST – Forced Swim Test. Mean±SEM; *P<0.05, **P<0.01

In adulthood (P56, **Fig. 2h**), ELA-exposed WT females showed significantly higher levels of immobility in the FST than standard-reared WT controls (**Fig. 2i,j**). This heightened immobility of ELA-exposed females was prevented by 5HT_1A_ K.D. (**Fig. 2i,j yellow line/bar**). Moreover, CORT responses to the FST were higher in ELA-exposed WT females compared to WT controls (**Fig. 2k**) and significantly reduced by 5HT_1A_ K.D. (**Fig. 2k, yellow line**). No significant effects of ELA or 5HT_1A_ K.D. on immobility or CORT were observed in adult males (**Fig. 2l-n**). When comparing baseline CORT levels between P35 and P56, CORT was significantly higher in adulthood at P56 in both sexes (**Supp. Fig. 2**).

Because we pooled mice from across different litters, we also test litter effects using “litter” as a covariate in our statistical analyses and found no effect of litter on the response to the FST (**Supp. Table 1**). LBN exposed WT and 5HT_1A_ K.D. offspring showed a reduction in body weight until P28, which recovered by P35 (**Supp. Fig. 3**).

In addition to ELA effects on acute stress responses to the FST, we also tested mice in the elevated plus maze (EPM) to assess innate avoidance behavior, which has been shown to be modulated by 5HT_1A_ autoreceptors. Consistent with previous studies,^50^ ELA did not affect avoidance of the open arms in WT male or female mice at P56. No effect of 5HT_1A_ K.D. was observed in males or females (**Supp. Fig. 4**).

Taken together, our data indicate that ELA in the form of LBN increases passive coping behavior and CORT responses to acute swim stress in adult females, which can be prevented by 5HT_1A_ autoreceptor knockdown.

### PVN activity and CRF protein levels are elevated by ELA and rescued by 5HT1A autoreceptor knockdown

A key step in initiating HPA axis responses and CORT release is the activation of neurons in the PVN of the hypothalamus. Because ELA increases CORT responses to an acute swim stress in adulthood, we next wanted to determine whether ELA also increases PVN activity and CRF levels. We, therefore, first used immunohistochemistry for the immediate early gene, cFos, as a proxy marker of neural activity in the PVN of ELA-exposed mice after acute exposure to FST stress in adulthood (**Fig. 3a-b**). ELA-exposed females showed higher numbers of cFos^+^ cells in the PVN compared to standard-reared controls, indicating higher levels of neural activity. This PVN hyperactivity was rescued by 5HT_1A_ K.D. (**Fig. 3c**), consistent with the regulation of CORT levels in these mice (as shown in **Fig. 2k**). ELA or 5HT_1A_ K.D. did not affect the number of cFos^+^ cells in the male PVN (**Fig. 3d**). Moreover, ELA-exposed females showed elevated levels of CRF protein in the PVN, which were rescued by 5HT_1A_ K.D. (**Fig. 3e**). No effects of ELA or 5HT_1A_ K.D. on CRF were detected in males (**Fig. 3f**).

**Figure 3:**
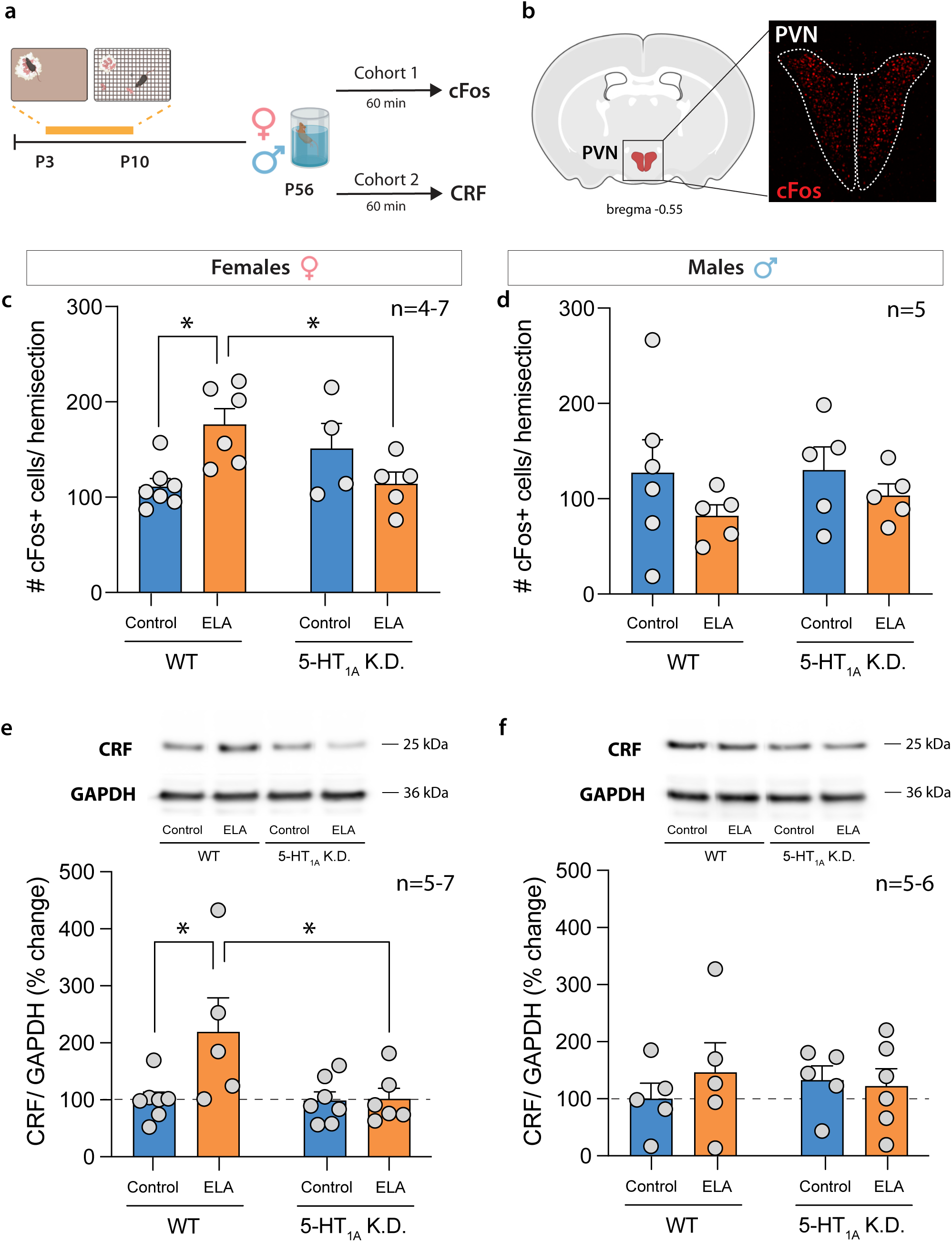
5-HT1A autoreceptor knockdown rescues PVN hyperactivity and CRF release in ELA-exposed females. **a.** Experimental timeline. **b.** Schematic and representative image of cFos+ cells in the PVN. **c.** ELA increases the number of cFos+ cells in the PVN of WT females compared to standard-reared control females. This effect was prevented in ELA-exposed 5-HT1A K.D. females. **d.** ELA or 5-HT1A K.D. did not affect cFos+ cells in the male PVN. **e.** ELA increased CRF in the PVN of WT females compared to controls. This effect was prevented in ELA-exposed 5-HT1A K.D. females. **f.** ELA or 5-HT1A K.D. did not affect CRF protein levels in the male PVN. Mean±SEM; *P<0.05

These data support the notion that ELA causes HPA axis hyperactivity at the level of the PVN in adult females, and that this effect can be prevented by 5HT_1A_ autoreceptor knockdown.

### Adult hippocampal neurogenesis is reduced following ELA in adulthood and rescued by 5HT_1A_ autoreceptor knockdown

Adult hippocampal neurogenesis is an important regulator of stress susceptibility^23^ and HPA axis activity.^24^ We, therefore, asked whether ELA may also reduce hippocampal neurogenesis as a potential mechanism predisposing to heightened stress vulnerability later in life. In adolescence (P35), we found no effect of ELA on the number of doublecortin (Dcx)-positive adult-born neurons in the dorsal or ventral DG of females or males (**Fig. 4a-d; Supp.** Fig 5a-d). However, Dcx^+^ adult-born neurons were reduced predominantly in the ventral DG of adult females with a history of ELA (**Fig. 4e; first two bars**), and to a lesser extent in the dorsal DG (**Supp.** Fig 5e). 5HT_1A_ K.D. counteracted the reduction in Dcx^+^ adult-born neurons in the vDG of ELA-exposed females (**Fig 4e; last two bars**). No effects of ELA or 5HT_1A_ K.D. were observed in males (**Fig 4f; Supp.** Fig 5f).

**Figure 4:**
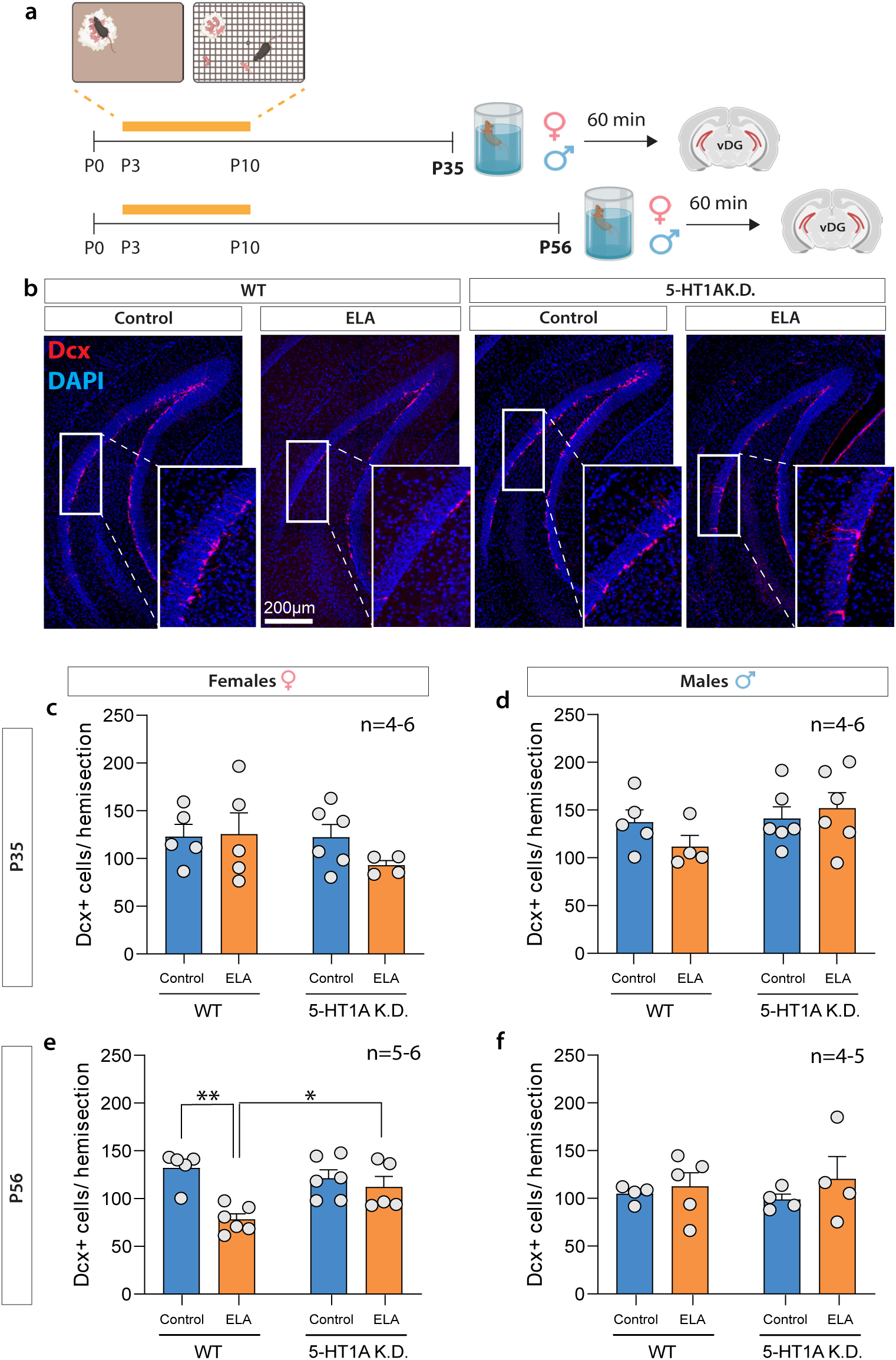
5-HT1A autoreceptor knockdown rescues ELA effects on adult hippocampal neurogenesis in the female vDG. **a.** Experimental timeline. **b.** Representative images of DCX^+^ cells in the female vDG at P56. **c.** In adolescence (P35), ELA or 5HT1A K.D. did not affect the number of DCX^+^ cells in the vDG of females, or **d.** males. **e.** In adulthood (P56), ELA decreased the number of DCX^+^ cells in the vDG of WT females compared to controls. This ELA effect was rescued in females with 5HT1A K.D. **f.** ELA or 5HT1A K.D. did not affect the number of DCX^+^ cells in the vDG of adult males. DCX – Doublecortin. Mean±SEM; *P<0.05; **P<0.01

Collectively, these data indicate that ELA reduces adult hippocampal neurogenesis predominantly in the vDG of adult females – a region of the DG that we previously identified to be crucial for stress vulnerability.^23^ Accordingly, this reduction in vDG neurogenesis is absent in 5HT_1A_ K.D. mice who do not exhibit enhanced stress reactivity in adulthood.

### Microglia activation is increased following ELA in adulthood and rescued by 5HT_1A_ autoreceptor knockdown

Previous studies have shown that ELA increases neuroinflammation.^42,43^ Inflammation also decreases hippocampal neurogenesis and increases behavioral and neuroendocrine responses to stress – effects of ELA that we see specifically in adult female mice. To test whether ELA and 5HT_1A_ K.D. cause sex-specific effects on neuroinflammation, we used immunohistochemistry for the microglia-specific calcium binding protein, Iba1, and the transmembrane glycoprotein, CD68, which is more abundant in phagocytic microglia following their activation in response to neuroinflammatory processes. We then quantified the number of CD68^+^/Iba1^+^ cells to determine microglia activation in the PVN and in the DG 1h after exposure to the acute swim stress (**Fig. 5a,b; Supp.** Fig 6a,b**; Supp.** Fig 7).

**Figure 5:**
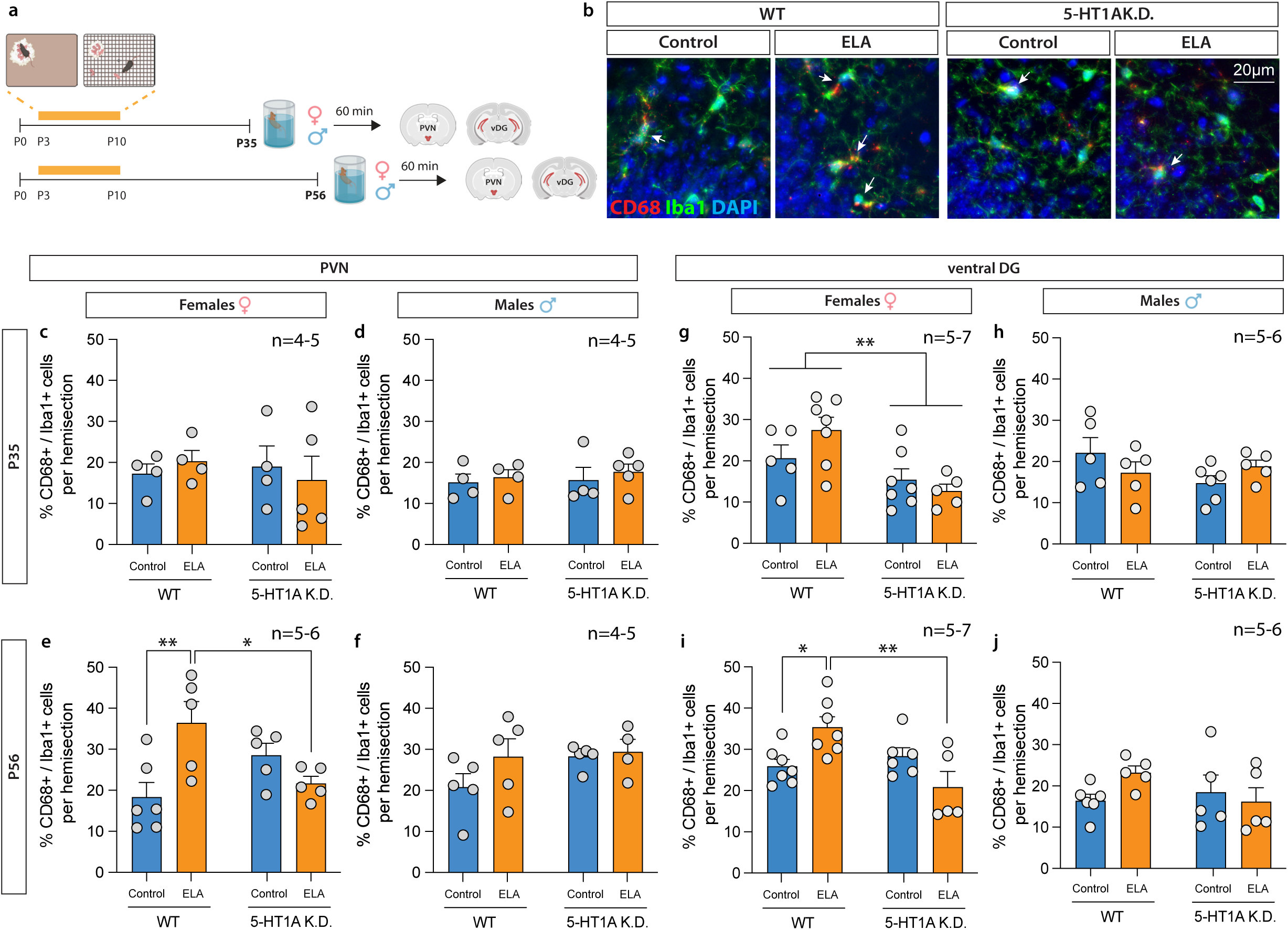
5-HT1A autoreceptor knockdown rescues ELA effects on microglia activation in the female PVN and vDG. **a.** Experimental timeline. **b.** Representative images of Iba1^+^ microglia (green) and CD68^+^ active microglia (red) in the female vDG. Arrowheads indicate CD68^+^/Iba1^+^ active microglia. **c-f.** Activated microglia in the PVN **c.** ELA or 5HT1A K.D. did not affect the % of active microglia in adolescent females or **d.** males. **e.** ELA increased the % active microglia in the PVN of adult WT females compared to controls. This effect of ELA was prevented in 5HT1A K.D. females. **f.** ELA or 5HT1A K.D. did not affect the % active microglia in the PVN of adult males. **g-j.** Activated microglia in the vDG. **g.** 5HT1A K.D. decreased the % active microglia in the vDG of adolescent females regardless of ELA exposure. **h.** ELA or 5HT1A K.D. did not affect the % of active microglia in adolescent males. **i.** ELA increased the % active microglia in the vDG of adult WT females compared to controls. This effect of ELA was prevented in 5HT1A K.D. females. **j.** ELA or 5HT1A K.D. did not affect the % active microglia in adult males. Mean±SEM; *P<0.05; **P<0.01

In adolescence (P35), we find no significant effect of ELA on the number of activated CD68^+^/Iba1^+^ microglia in the PVN (**Fig. 5c,d**), ventral DG (**Fig. 5g,h**) or dorsal DG (**Supp. Fig. 6**c,d) of males and females. However, 5HT_1A_ autoreceptor knockdown reduced overall microglia activation in the vDG of adolescent females (**Fig. 5g**). In adulthood (P56), microglia activation was increased in the PVN (**Fig. 5e**) and in the ventral DG of ELA-exposed WT females (**Fig. 5i**), and this effect was rescued by 5HT_1A_ K.D. (**Fig. 5e,i, last column**). No effects of ELA or 5HT_1A_ knockdown were observed in the PVN and ventral DG of males (**Fig. 5f,j**) or in the dorsal DG of either males or females at P56 (**Supp. Fig. 6**e,f).

These data show that ELA causes sex-dimorphic effects on neuroinflammation in the PVN and ventral DG that manifest predominantly in adult female mice and that can be prevented by 5HT_1A_ autoreceptor knockdown.

## DISCUSSION

Identifying neurobiological mechanisms that can prevent ELA effects on future stress reactivity may help develop novel strategies to treat or prevent psychiatric disorders that have their origin early in life. Here, we show that ELA in the form of LBN causes persistent sex-dependent impairments in 5HT neurons and in behavioral and neurobiological responses to stress. We find that these stress vulnerability-promoting effects of ELA are specific to females, develop primarily in adulthood, and can be prevented by postnatal knockdown of 5HT_1A_ autoreceptors. Our findings shed new light on the biological mechanisms that promote resilience to ELA on stress response regulation across development and highlight the 5HT_1A_ receptor as a potential target for intervention.

To determine how ELA affects stress reactivity later in life, we raised mice under LBN conditions and exposed them to an acute FST as a means by which to induce an acute stress response later in life. We find that in females, ELA increases immobility in the FST, possibly indicating a more passive approach to coping with the inescapable swim stress. This heightened behavioral response to stress of ELA-exposed females emerges in young adulthood (at P56), but not earlier in adolescence (at P35) (**Fig. 2**). Accordingly, we find heightened neuroendocrine responses to stress in ELA-exposed females, evidenced by higher CORT responses to the FST, higher PVN activity, and more CRF protein levels in PVN tissue of ELA-exposed females compared to standard-reared controls (**Figs. 2,3**). While we investigated PVN activity by quantifying the number of cFos^+^ cells in the entire PVN, it is important to note that the PVN contains different neuron subtypes in addition to CRF neurons, including, e.g., oxytocin and vasopressin neurons. Future studies specifically investigating ELA effects on the activation of CRF neurons or other neural subtypes in the PVN will be important to further characterize how early experiences affect PVN regulation of these different neuroendocrine systems. While increased PVN cFos and CRF levels together with heighted serum CORT levels following ELA support the notion of HPA axis hyperactivity, a caveat of measuring CRF protein in PVN tissue micropunches is that higher CRF levels could reflect either increased production and turnover, or reduced secretion. It is also noteworthy that while baseline CORT levels of ∼200 ng/ml in our study are consistent with previous studies (e.g.,^63,64,65^) they are higher than the more commonly reported ∼50 ng/ml. In our experiments, all mice were transferred and acclimated to a new room 1h prior to blood collection, which may have potentially increased baseline CORT release in the home cage due to transport out of the housing room.

We also find that ELA decreases neurogenesis in the vDG of adult females and increases neuroinflammation in the PVN and vDG, but not in the dDG. Reduced adult hippocampal neurogenesis and increased neuroinflammation specifically in the vDG have previously been associated with heightened stress vulnerability,^5,20,21,23,66–68^ and both hippocampus and PVN have been implicated in coping behavior and immobility in the FST.^69,70^ Higher inflammation and reduced neurogenesis in these regions may thus contribute to the neuroendocrine and behavioral effects of ELA in our experiments.

We find that the effects of ELA on stress reactivity, neurogenesis, and neuroinflammation, are absent in mice with postnatal knockdown of raphé 5HT_1A_ autoreceptors. Under physiological conditions, 5HT_1A_ autoreceptors inhibit raphé 5HT neurons and 5HT release at their terminal projection sites. Indeed, we find that ELA reduces expression of the immediate early gene, cFos, in raphé TPH2^+^ cells of adult females, indicating that reduced activity of 5HT neurons following ELA primarily emerges in adulthood (**Fig. 1**). This adult impairment in 5HT neuron activity may thus be underlying the behavioral and neurobiological deficits following ELA, which also manifest predominantly in adulthood and are rescued by disinhibiting 5HT neurons through 5HT_1A_ autoreceptor knockdown. This conclusion is supported by previous work using the same transgenic mouse model and showing that 5HT_1A_ knockdown in adulthood increases spontaneous firing of 5HT neurons and reduces vulnerability to an acute FST stress in adult male mice.^56^ While we quantified cFos as a proxy marker for neural activity, future studies should be aimed at investigating how ELA affects the intricate electrophysiological properties of 5HT neurons that cause these changes in cFos expression. In our study, 5HT_1A_ autoreceptor knockdown was induced perinatally and maintained until adulthood. Resilience to the effects of ELA in our experiments is therefore likely a result of the combined 5HT_1A_ autoreceptor reduction during ELA and during adolescent and adult development. Disentangling the impact of 5HT_1A_ as a resilience promoting factor during the experience of ELA vs a treatment target after ELA will be important to further refine the possible therapeutic potential of this receptor and of disinhibiting 5HT neurons. While we find no effect of ELA or 5HT_1A_ knockdown on innate avoidance, previous studies have indeed shown that either adult or juvenile 5HT_1A_ autoreceptor knockdown increase avoidance.^62,71^ It is possible that continuous postnatal knockdown of 5HT_1A_ autoreceptors, as used in our study, may have different effects on avoidance than autoreceptor knockdown during discrete developmental periods or germline knockdown of 5HT_1A_ auto- and heteroreceptors, possibly due to interactions with other developmental processes. While the purpose of our study was to elucidate ELA effects on stress reactivity, it is possible that 5HT_1A_ knockdown may affect behavior in other avoidance tasks.

The pronounced sex-specific effects of ELA in our study are consistent with previous findings showing heightened vulnerability of females to the behavioral consequences of LBN.^50,51^ It is worth noting, however, that LBN-reared male offspring show similar reductions in body weight as females until P35, suggesting that LBN conditions do indeed have some effects also on males (**Supp. Fig. 3**c). Accordingly, previous studies have shown that LBN causes behavioral and biological impairments in male mice, including altered spatial recognition abilities,^58^ feeding behavior,^72^ and mitochondria function.^73^ It is thus possible that males are not simply more resilient to the effects of ELA, but that ELA may affect different neurobiological mechanisms and their associated behavioral outcomes in male vs female mice. Previous studies have suggested sex- and sex-hormone dependent regulation of glucocorticoid receptor (GR) function in the PVN.^74,75,76^ It is possible that sex-specific GR expression could cause differences in HPA axis responsiveness to ELA, which in turn may lead to sex-specific effects of ELA on stress responsivity in adulthood, as we observed in our study. This possible mechanism should be investigated by future studies.

The 5HT_1A_ receptor has been implicated in the pathogenesis of anxiety disorders, major depression, suicidal ideation, and in the mechanism of action of antidepressants.^61,70,76–88^ Indeed, new compounds targeting 5HT_1A_ have emerged over recent years that are currently being used or tested for their antidepressant- or anxiolytic effects.^89–92^ Our data support the notion that a developmental reduction in the function of presynaptic 5HT_1A_ autoreceptors in the raphé nuclei prevents ELA effects on future stress sensitivity. While chronic SSRI treatment in adulthood is known to downregulate 5HT_1A_ autoreceptors, the timing of 5HT_1A_ modulation during development may be critical in establishing long-lasting resilience to stress. Targeting 5HT_1A_ autoreceptors with early life interventions could potentially rewire stress-responsive neural networks, in turn providing long-lasting resilience to future stressors. However, the use of SSRIs in children raises important safety concerns, underscoring the need to discover novel compounds targeting 5HT_1A_ and to improve our knowledge about affected neural circuits. We have previously shown that activating postsynaptic 5HT_1A_ heteroreceptors in the DG promotes resilience to chronic stress,^52^ in addition to their reported role in mediating antidepressant effects on behavior.^88^ Recent advances in the development of biased 5HT_1A_ ant/agonists with functional selectivity at either 5HT_1A_ autoreceptors or 5HT_1A_ heteroreceptors may thus enable harnessing the differential potential of these receptors in specific cell types and brain regions in the future.^93^ Identifying novel drugs with pharmacological profiles that can target specific subsets of 5HT_1A_ receptors or their downstream signaling pathways will be particularly important for patients with a history of ELA who respond poorly to currently available antidepressants.^94^ Preventing the enduring consequences of ELA on 5HT neuron function in adulthood, as accomplished in our study by developmental knockdown of 5HT_1A_ autoreceptors, could help alleviate heightened stress vulnerability in individuals with a history of ELA.

## ACKNOWLEDGEMENTS

We would like to thank Dr. Eduardo Dave Leonardo for providing the Pet1-tTS;tetO-1A mice. We thank Dr. Maryam Hasantash for help with covariate analyses, and Ms. Baoyao Huang for technical assistance with RNAscope experiments.

## AUTHOR CONTRIBUTIONS

RSD, LM, RT, SW, YL, NS, and CA, contributed to data acquisition, analysis, and interpretation. CA designed and drafted the work. CA, RT, and YL contributed to revising the work.

## FUNDING

This project was supported by grants from the National Institute of Mental Health (NIMH) (R00MH108719, P50MH090964, P50MH090964-S2, R01MH126105), a NARSAD Young Investigator Award (to C.A), and by a Columbia University Institute for Developmental Sciences Award (to L.M.).

## DATA AVAILABILITY STATEMENT

The datasets generated during and/or analyzed during the current study are available from the corresponding author on reasonable request.

## DISCLOSURES

Dr. Anacker has received research funding from Sunovion Pharmaceuticals and consulting fees from Ono Pharmaceuticals. All other authors have nothing to disclose.

## Notes

### Competing Interest Statement

Dr. Anacker has received research funding from Sunovion Pharmaceuticals and consulting fees from Ono Pharmaceuticals unrelated to the work in this publication. All other authors have nothing to disclose.

### Summary of Updates

New data has been added in Figure 1, showing the extent of the 5HT1A knockdown across development. New data has been added showing the effects of ELA and 5HT1A knockdown on activity of raphe serotonin neurons. All Figures have been updated to show individual data points.

