## Supplemental Materials for "Sex-specific and Developmental Effects of Early Life Adversity on Stress Reactivity are Rescued by Postnatal Knockdown of 5-HT_1A_ Autoreceptors"

*Dixon, Malave, et al.*

### SUPPLEMENTAL METHODS

#### Experimental Setup

All experimental procedures were conducted in accordance with the US National Institutes of Health (NIH) Guide for the Care and Use of Laboratory Animals, the New York State Psychiatric Institute (NYSPI) Institutional Animal Care and Use Committee (IACUC). All experimental procedures were conducted in accordance with the US National Institutes of Health (NIH) Guide for the Care and Use of Laboratory Animals, the New York State Psychiatric Institute (NYSPI) Institutional Animal Care and Use Committee (IACUC) at Columbia University, and the Research Foundation for Mental Hygiene (RFMH). Pet1-tTS;tetO-1A mice homozygote for the tetracycline operator (tetO) in the promoter region of the *5htr1a* gene and heterozygote for the tetracycline-dependent transcriptional suppressor (tTS) under control of the 540Z Pet-1 promoter fragment (Pet1-tTS)[59] were used for all experiments. In the presence of doxycycline (DOX), raphé $5HT_{1A}$ levels are indistinguishable between mice heterozygote for the *Pet1-tTS* transgene (Pet1-$tTS^+$) and mice homozygote negative for the *Pet1-tTS* transgene (Pet1-$tTS^-$). Removal of DOX causes suppression of *5htr1a* expression in transgenic heterozygote Pet1-$tTS^+$ mice (from here on referred to as 5-HT1A K.D.) but not in homozygote negative Pet1-$tTS^-$ mice (from here on referred to as WT).[59] Same sex mice were housed 3–5 per cage with *ad libitum* access to food and water on a 12:12h light/dark cycle. Nulliparous Pet1-$tTS^-$ females were bred with Pet1-$tTS^+$ males at P56 and maintained on DOX diet. DOX was removed three days before parturition. Breeding Pet1-$tTS^-$ females with heterozygote Pet1-$tTS^+$ males results in litters with 50% heterozygote Pet1-$tTS^+$ offspring (5-HT1A K.D.) and 50% Pet1-$tTS^-$ offspring (WT). Female and male Pet1-$tTS^-$ (WT) and Pet1-$tTS^+$ (5-HT1A K.D.) offspring were then raised on regular chow and weaned at P21. Separate cohorts of mice were used for behavior testing in the Forced Swim Test (FST) at P35 and P56, and in the Elevated Plus Maze (EPM) at P56. Behavioral testing was conducted during the light period between 10 am and 2 pm. Of all mice that underwent the FST a subset of mice was perfused with 4% paraformaldehyde (PFA) for immunohistochemistry of cFos in the PVN, doublecortin (Dcx) in the dorsal and ventral dentate gyrus (DG), and CD68+/Iba1+ activated microglia in the PVN, dorsal DG, and ventral DG. Another subset of mice exposed to the FST was subjected to live decapitation and brains flash frozen in ice cold isopentane for CRF protein analysis by Western Blot. To assess 5HT1A K.D. by RNAscope, additional separate cohorts of mice were live decapitated at P3, P10, P35, and P56, and brains flash frozen in ice cold isopentane.

**Forced Swim Test (FST)**

We used the Forced Swim Test (FST) to induce acute stress in adolescence (P35) and adulthood (P56) as previously described.[1] Briefly, mice were placed in individual cylinders (46cm tall x 32cm in diameter) containing 25-26°C water (room temperature) 30cm deep for 6 minutes. Swim sessions were recorded from a video camera directly above the swim buckets. Following swim sessions, mice were removed from the cylinders, dried with paper towels, placed in a cage over a heating pad for 15 min, and then returned to their home cages. Swim sessions were analyzed using the automated tracking system VideoTrack software (ViewPoint). Immobility was measured when the animal exhibited no movement or floated in the water without struggle. Time spent immobile was reported in seconds during one-minute bouts.

**Elevated Plus Maze (EPM)**

We used the Elevated Plus Maze (EPM) to test avoidance behavior in adult female and male mice at P56, as previously described.[1] Briefly, the EPM is a plus-cross-shaped apparatus consisting of four arms, two open and two enclosed by walls, linked by a central platform at a height of 50 cm from the floor. Mice were individually placed in the center of the maze facing an open arm and were allowed to explore the maze for 5 min. The time spent in the open arms was used as an index of avoidance behavior. Videos were scored using ANY-maze behavior tracking software (Stoelting, Wood Dale, IL).

**Corticosterone (CORT) EIA assay**

Submandibular blood draw was done using Goldenrod animal lancets (Braintree Scientific, Inc) 30 min after mice were exposed to the FST. For baseline CORT assessment, mice were transferred and acclimated to a new blood collection room 1h prior to submandibular blood draw. Blood was centrifuged at 3000 rpm for 15 min at 4°C and the upper serum fraction was collected stored at -80°C. Corticosterone EIA assays (ArborAssays, Ann Arbor, MI) were run according to the manufacturer's instructions. Briefly, 5 µl of serum were diluted 1:150 in Assay Buffer and added to the assay plate. DetectX Corticosterone Conjugate (25 µl) and DetectX Corticosterone antibody (25 µl) were added to each well and incubated on a shaker for 1 h at room temperature. The assay plate was washed 4 times with 300 µl of wash buffer and 100 µl of TMB substrate were added to each well for 30 min without shaking. Reactions were stopped with 50 µl HCl-based Stop buffer and optical density was read at 450 nm. Sample absorbance values were compared to a standard curve on the same assay plate, ranging from 78,15 pg/ml to 10,000 pg/ml.

**Immunohistochemistry**

Free-floating brain sections containing PVN, dorsal hippocampus, and ventral hippocampus were washed in PBS + 0.3% Triton X-100 (PBST), incubated in 10% normal donkey serum (NDS) in PBST for 2h, and incubated overnight at 4 °C with primary antibody in 10% NDS/PBST (rabbit-anti-cFos, 1:2000, SySy; rabbit-anti-Dcx, 1:1500, Cell Signaling; rabbit-anti-iba1, 1:1500, Wako Chemicals; Rat-anti-CD68, 1:1000, Bio-Rad). The next day, sections were washed with 1x PBST three times for 10 min and incubated in secondary antibody (donkey anti-rabbit IgG Alexa 594, 1:500; donkey anti-rabbit IgG Alexa 488, 1:500; donkey anti-rat IgG Alexa Cy5, 1:500, Jackson ImmunoResearch) in 10% NDS/PBST for 2h at room temperature, washed with 1xPBS twice for 10 min, and incubated with 1xPBS + DAPI (1:5,000; Sigma-Aldrich) for nucleic acid staining for 10 min. Sections were mounted onto glass slides with AquaPolymount anti-fade mounting medium and imaged on a Leica fluorescent microscope.

**RNAscope**

To detect TPH2, 5HT1A, and cFos mRNAs in coronal brain slices, RNAscope was performed using Multiplex Fluorescent V2 Assay (Advanced Cell Diagnostics, ACD) on frozen tissue samples according to the manufacturer's protocol. All steps were carried out at room temperature unless stated otherwise. Mice were live decapitated and brains flash frozen in ice-cold isopentane. Brains were sectioned at −20°C in 16 µm-thick coronal slices using a cryostat (Leica CM 3050S) and directly mounted onto glass slides. Sections were incubated with 4% PFA for 1.5h, rinsed with PBS, dehydrated in increasing concentrations of ethanol (50%, 70%, 100%, 5 min each), air-dried, treated with hydrogen peroxide for 10 min, rinsed with dH2O, baked for 30 min at 37°C in an HybEZ II oven (ACD 321721), and treated with Protease IV for 30 min at RT and washed with dH2O. Probes were hybridized for 2hrs at 40°C. Sections were then incubated with AMP1 for 30 min at 40°C, AMP2 for 30 min at 40°C, and AMP3 for 15 min at 40°C in the oven. RNA hybridization antisense probes were consecutively developed with VIVID 570 dye (5HT1A-C2), VIVID 650 dye (cFos-C3), and VIVID 520 dye (TPH2-C1), counterstained with DAPI, air dried, coverslipped, and stored at 4°C.

**Fluorescent microscopy**

To identify fluorescently labeled cells, images of raphé, PVN, dorsal DG, and ventral DG were taken using a Leica Thunder Tissue Imager (Leica DM6 B, Leica Microsystems Inc.) equipped with LAS X Navigator Lightning Software. Immunofluorescent sections were excited at 405nm

(DAPI), 488nm (Iba1), 594nm (cFos, DCX), and 647nm (CD68). RNAscope fluorophore labeled sections were excited at 488nm (TPH2), 590nm (5HT1A), and 605nm (cFos). cFos and Dcx immunofluorescent sections were imaged with a Leica 20x dry objective (numerical aperture 0.80, working distance 0.4 mm), with a FOV of 553.6 x 553.6 μm, a pixel size of 0.5 x 0.5 μm, and a speed of 400 Hz. Immunofluorescent CD68/Iba1 sections and RNAscope fluorophore labeled 5HT1A/TPH2 and cFos/TPH2 sections were imaged with a Leica 40x dry objective (numerical aperture 0.95, working distance 0.17mm) with a FOV of 333.7 × 333.7 μm, a pixel size of 0.16 × 0.16 μm, and a speed of 40 Hz. Z-stack images were taken with a step size of 0.30 μm. Equal exposure times were used for all images (RNAscope: 5HT1A: 28ms, cFos: 60ms, TPH2: 3.4ms; Immunohistochemistry: cFos: 180ms, DCX: 340ms, CD68: 167ms, Iba1: 160ms, DAPI: 2.7ms).

**Cell Quantification**

To quantify fluorescently labeled RNAscope images of 5HT1A, cFos, and TPH2 mRNA expression in raphé neurons, we used a standardized quantification pipeline in QuPath0.5.1. Cell nuclei were identified based on DAPI fluorescence and cells were defined with 5μm sigma and 7μm cell expansion around each cell nucleus in QuPath. To determine % $5HT1A^+/TPH2^+$ and % $cFos^+/TPH2^+$ cells, composite classifiers were trained in QuPath. For each classifier, grayscale images of each individual fluorescent channel were thresholded based on the following cell mean pixel intensity thresholds: 5HT1A: 100; cFos: 60; TPH2: 300. Thresholds were consistently applied across all images and all experimental conditions. To quantify the % of raphé serotonin neurons expressing 5HT1A, the number of 5HT1A/TPH2 double-positive cells was plotted as a percentage of the total number of $TPH2^+$ cells. To quantify the % of active raphé serotonin neurons, the number of cFos/TPH2 double-positive cells was plotted as a percentage of the total number of $TPH2^+$ cells.

To quantify immunofluorescent images of cFos and Dcx, both hemispheres of each mouse were counted on every sixth section along the longitudinal axis of the hippocampus (4-8 sections in total). Cell counts from each hemisphere were averaged across one section and then each section was averaged for the dorsal DG or ventral DG region for each mouse. For quantification of CD68/Iba1, the tip, superior blade and inferior blade of the DG was imaged at 40x for both hemispheres of each mouse along the longitudinal axis of the hippocampus as described above. To quantify active microglia, the number of active microglia ($CD68^+$ and $Iba1^+$ cells) was plotted as a percentage over the total number of microglia ($Iba1^+$ cells). All images were counted by an experimenter blind to experimental condition.

**Western Blot Analysis**

To obtain PVN protein samples, mice were live decapitated, and brains flash frozen in ice cold isopentane and stored at -80°C. Transversal brain sections (200 µm) were obtained as previously described[2] and PVN tissue biopsies collected. Tissue biopsies were incubated in 150ul Neuronal Protein Extraction Reagent (N-PER™, Thermo Scientific) for 30 min and vortexed rigorously every 10min. Protein concentrations were quantified using a bicinchoninic acid (BCA) colorimetric assay system (Pierce). Protein samples containing 20 µg of total protein were boiled for 10 min at 95 °C in NuPAGE LDS sample buffer (Invitrogen) and NuPAGE sample reducing agent (Invitrogen) and subjected to reducing SDS/PAGE on 10% NuPAGE Bis-Tris gels for 35min at 200 V. Proteins were electrophoretically transferred to Immuno-Blot PVDF membranes (Bio-Rad) at 35 V for 1.5 h at 4 °C. Transfer efficiency was confirmed using pre-stained protein standards. Unspecific binding sites were blocked for 1 h in 5% nonfat dry milk in PBS and membranes were immunoprobed with polyclonal rabbit anti-CRF (1:1,000, Abcam, ab184238) and rabbit anti-Glyceraldehyde 3-phosphate dehydrogenase (GAPDH) antibody (1:20,000, Proteintech, 10494-1-AP) in blocking solution at 4 °C overnight. Membranes were washed three times for 10min with PBS containing 0.1% Tween 20 (PBST) and incubated with goat anti-rabbit HRP secondary antibody (1:10,000, Jackson Immunoresearch) in 5% nonfat dry milk in PBS for 1 h at room temperature. Membranes were washed in PBST and imaged using Azure c600 imaging station. CRF protein levels were normalized to GAPDH protein levels in each sample and differences in normalized CRF/GAPDH levels are expressed as % of WT control levels.

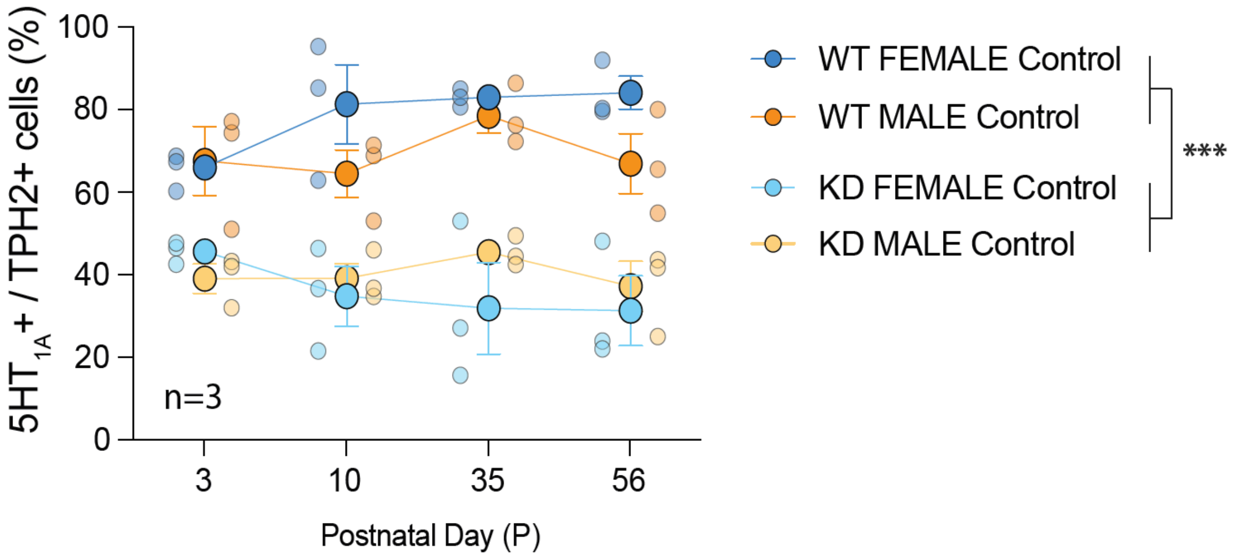


**Supplementary Figure S1: Sex does not affect $5HT_{1A}$ K.D.** RNAscope data of $5HT_{1A}$ mRNA expression in $TPH2^{+}$ raphe 5HT neurons of male and female WT and $5HT_{1A}$ K.D. mice at postnatal days (P) 3, 10, 35, and 56. Mean±SEM; ***P<0.001 indicates main effect of genotype (knockdown).

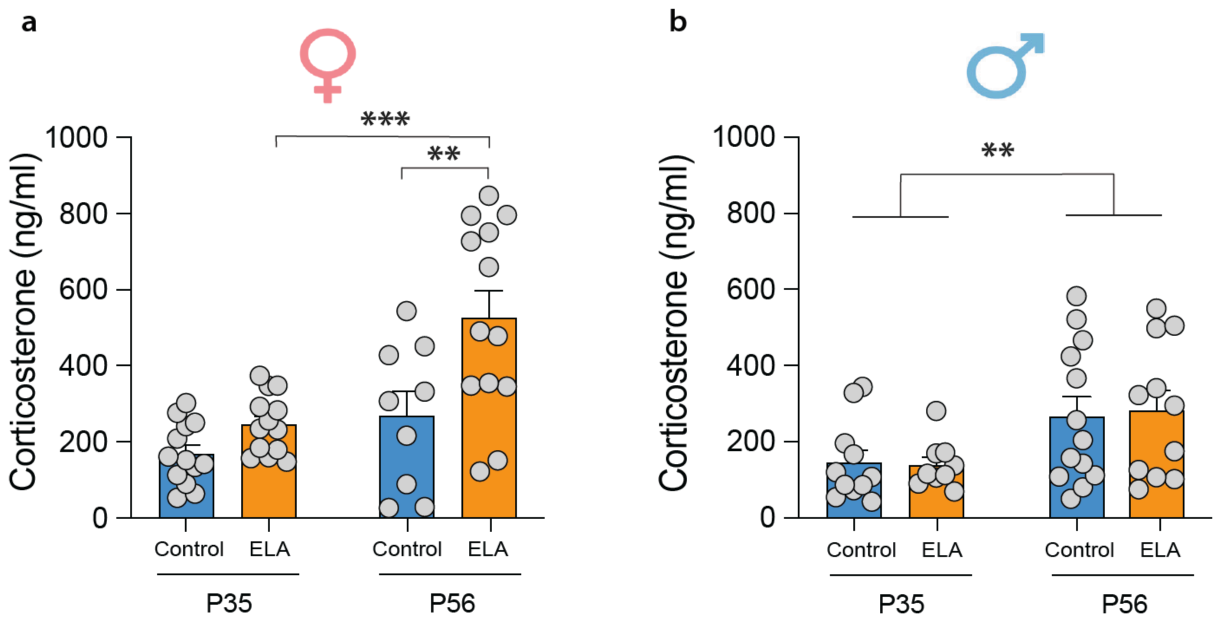


**Supplementary Figure S2: Baseline CORT comparisons across age. a.** Female baseline CORT levels at P35 and P56 in WT mice. **P<0.01, ***P<0.001 denote posthoc tests. **b.** Male baseline CORT levels at P35 and P56 in WT mice. **P<0.01 denotes main effect of age.

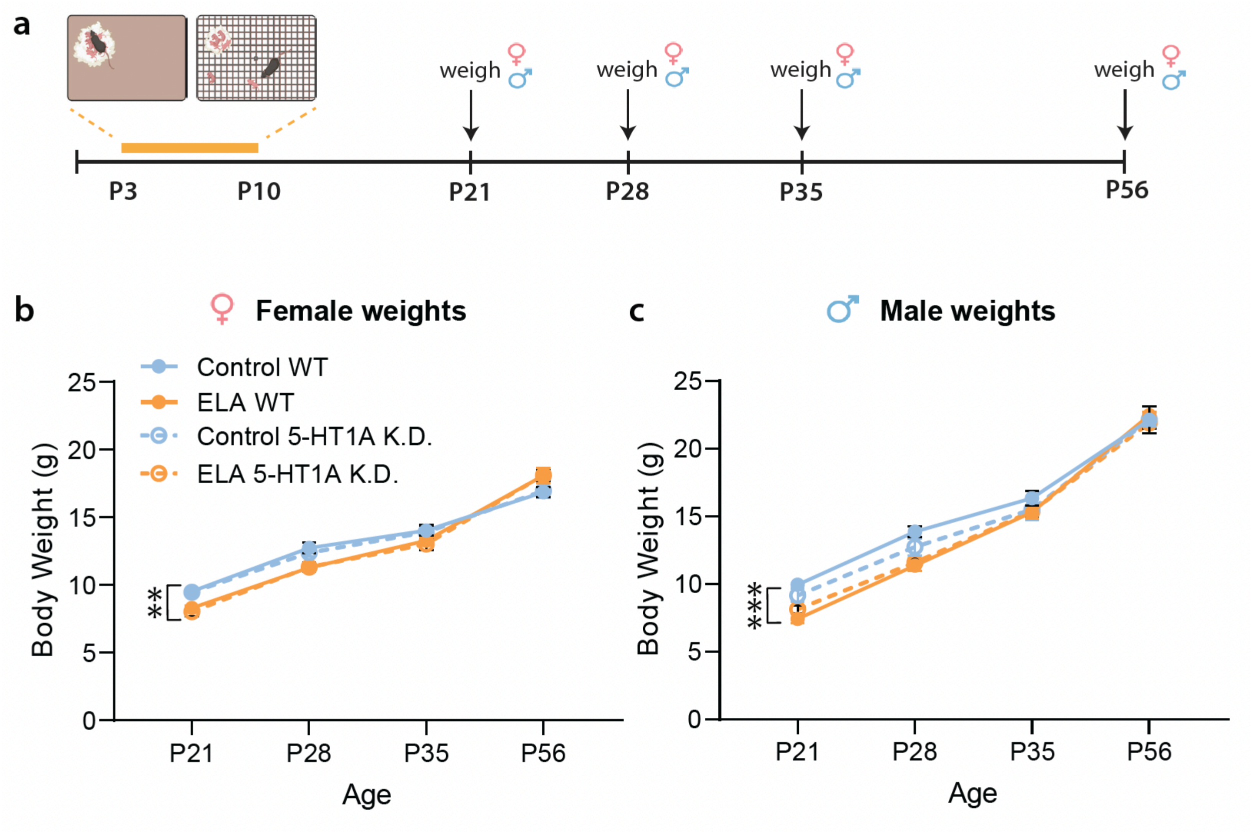


**Supplementary Figure S3: ELA reduces body weight until P35. a.** Timeline of LBN model and body weight measurements. **b.** Body weight of female mice at age P21, P28, P35 and P56. ELA reduces body weight of both WT and 5HT1A K.D. females compared to controls until P35. **c.** Body weight of male mice at age P21, P28, P35 and P56. ELA reduces body weight of both WT and 5-HT1A K.D. males compared to controls until P35. Mean±SEM; **P<0.01, ***P<0.001 indicate main effect of ELA.

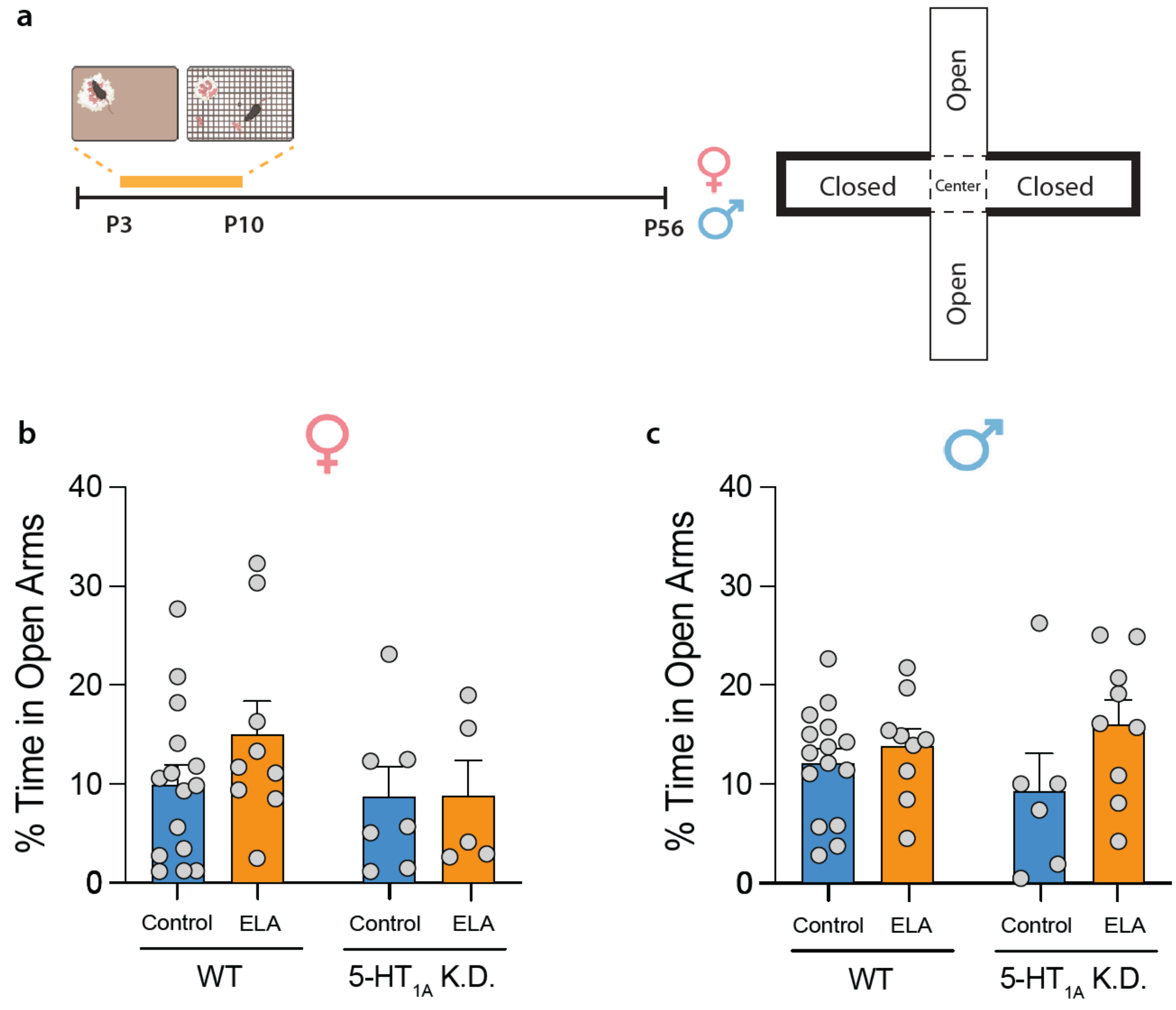


**Supplementary Figure S4**: **No effect of ELA or 5HT1A K.D. on open arm avoidance. a.** Timeline of ELA exposure and testing in the elevated plus maze (EPM) at P56. **b.** No effect of ELA or 5HT1A K.D. on open arm avoidance in females or **c.** males.

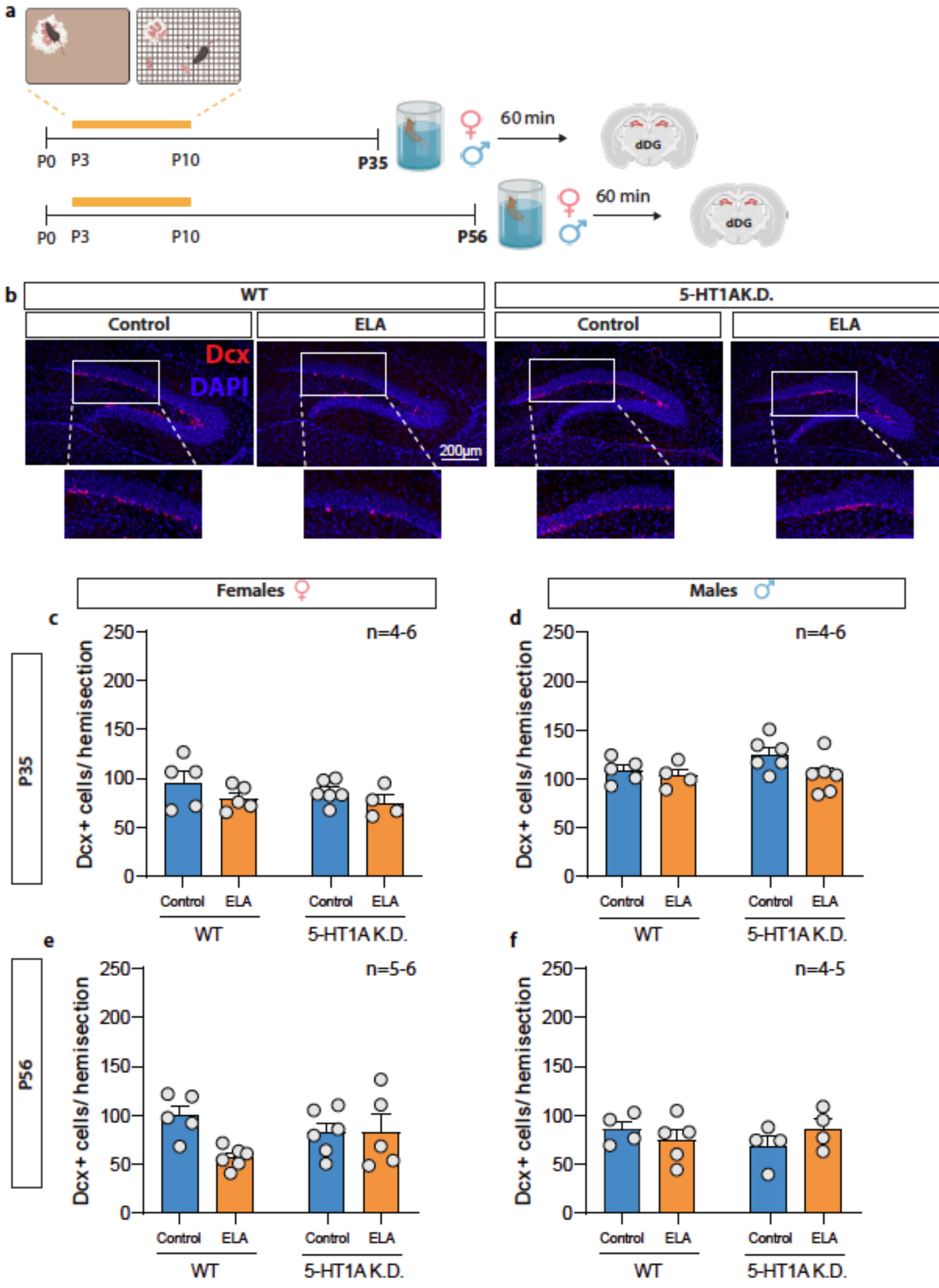


**Supplementary Figure S5**: **No effect of ELA or 5HT1A K.D. on adult hippocampal neurogenesis in the dDG. a.** Timeline of LBN model and tissue collection. **b.** Representative images of DCX$^{+}$ cells in the female dDG. **c.** In adolescence (P35), neither ELA nor 5-HT1A K.D. affected the number of DCX$^{+}$ cells in the dDG of females, or **d.** males. **e.** In adulthood (P56), no significant effects of ELA or 5-HT1A K.D. were observed on the number of DCX$^{+}$ cells in the dDG females, or **f.** males. DCX – Doublecortin. Mean±SEM; n=4-6

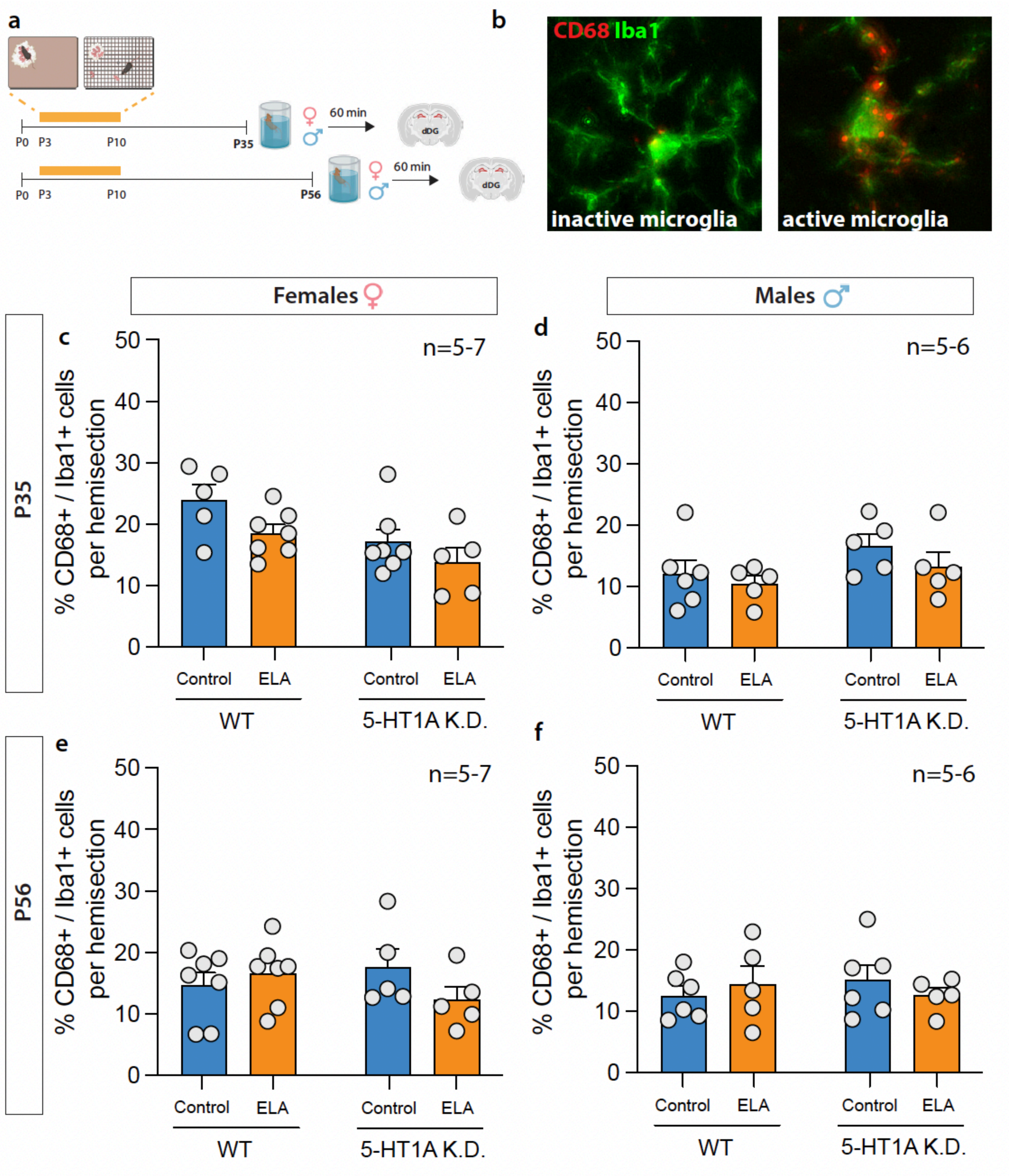


**Supplementary Figure S6**: **No effect of ELA or 5HT1A K.D. on microglia activation in the dDG. a.** Timeline of LBN model and tissue collection. **b.** Representative images of Iba1$^{+}$ cells (green) and CD68$^{+}$ cells (red) showing inactive Iba1$^{+}$ microglia (left panel) and active CD68$^{+}$/Iba1$^{+}$ microglia (right panel). **c.** In adolescence (P35), neither ELA nor 5-HT1A K.D. affected the number of CD68$^{+}$/Iba1$^{+}$ active microglia in the dDG of females, or **d.** males. **e.** In adulthood (P56), no significant effects of ELA or 5-HT1A K.D. were observed on the number of CD68$^+$/Iba1$^+$ active microglia in the dDG of females, or **f.** males. Mean±SEM; n=5-7

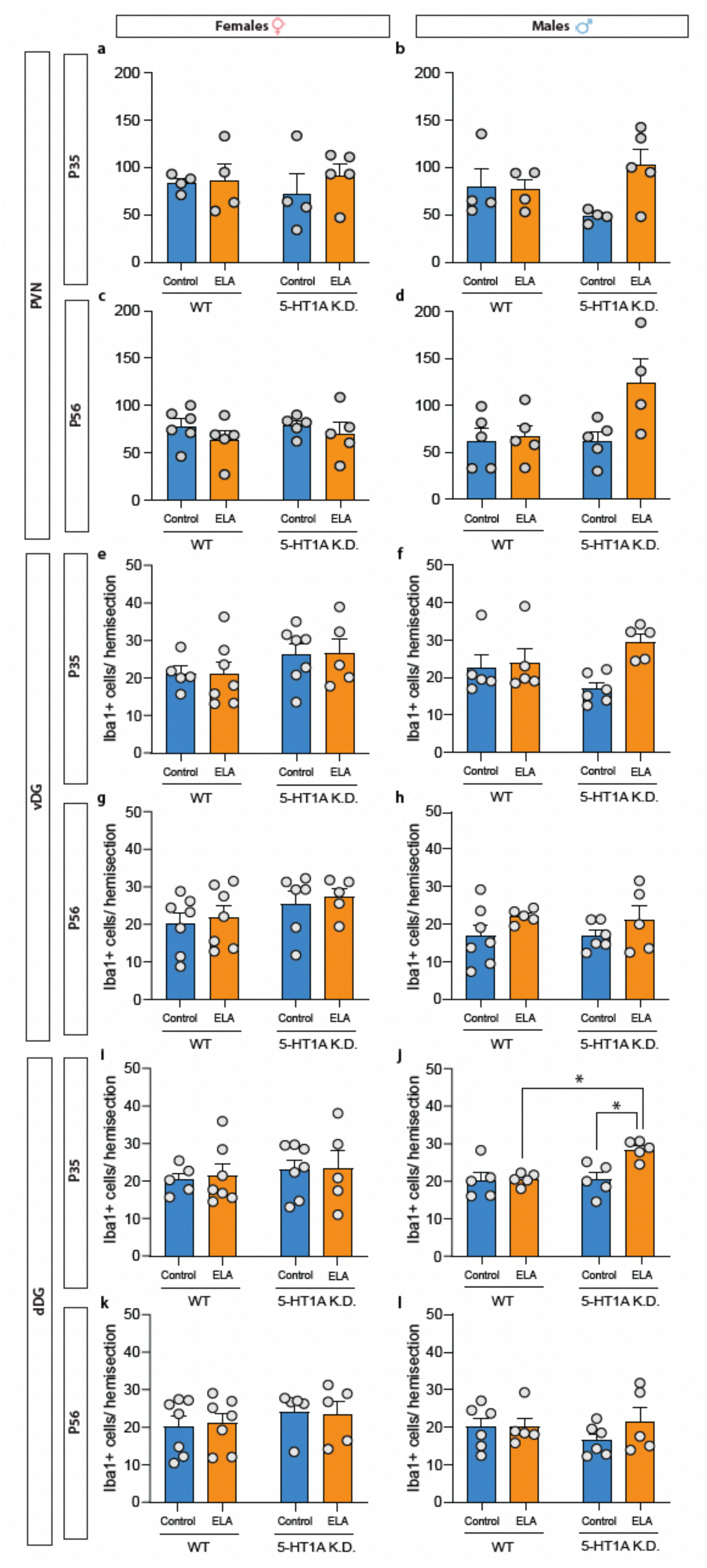
Females ♀
Males ♂
PVN
vDG
dDG
P35
P56
a
b
c
d
e
f
g
h
i
j
k
l
Iba1+ cells/ hemisection
Control
ELA
WT
5-HT1A K.D.
*

**Supplementary Figure S7**: **ELA and 5-HT1A K.D. effects on total numbers of Iba1$^{+}$ microglia in the PVN, vDG and dDG. a.** No significant group differences were observed in the total number of Iba1$^{+}$ microglia in the PVN of adolescent females or **b.** males. **c.** No significant group differences were observed in the total number of microglia in the PVN of adult females or **d.** males. **e.** No significant group differences were observed in the total number of Iba1$^{+}$ microglia in the vDG of adolescent females or **f.** males. **g.** No significant group differences were observed in the total number of microglia in the vDG of adult females or **h.** males. **i.** No group differences in the total number of microglia in the dDG of adolescent females. **j.** 5-HT1A K.D. increased the total number of microglia in ELA-exposed males in adolescence. **k.** No group differences were observed in the total number of microglia in adult females or **l.** males. Mean±SEM; n=5-7; *P<0.05 indicates Tukey *post-hoc* test.

### Supplementary Table 1. Detailed Statistics

| Name | Abbrev | Measurement | Statistical Test | Comparison | F | ° of | p | * | Fig. | Post hoc Test |
|---|---|---|---|---|---|---|---|---|---|---|
| Figure 1 5-HT effects | 5HT1A K.D. | Females Ctrl WT vs Ctrl K.D. | 2-Way ANOVA | Interaction | 2.54 | 3, 16 | 0.09 | ns | 1d | n.a. |
| | | | | Time | 0.04 | 3, 16 | 0.99 | ns | | |
| | | | | GT | 80.37 | 1, 16 | <0.0001 | *** | | |
| | | Males Ctrl WT vs Ctrl K.D. | 2-Way ANOVA | Interaction | 0.18 | 3, 16 | 0.91 | ns | 1e | n.a. |
| | | | | Time | 1.55 | 3, 16 | 0.24 | ns | | |
| | | | | GT | 55.78 | 1, 16 | <0.0001 | *** | | |
| | | Females ELA WT vs ELA K.D. | 2-Way ANOVA | Interaction | 0.46 | 2, 12 | 0.64 | ns | 1d | n.a. |
| | | | | Time | 0.17 | 2, 12 | 0.84 | ns | | |
| | | | | GT | 45.24 | 1, 12 | <0.0001 | *** | | |
| | | Males ELA WT vs ELA K.D. | 2-Way ANOVA | Interaction | 0.49 | 2, 12 | 0.63 | ns | 1e | n.a. |
| | | | | Time | 3.11 | 2, 12 | 0.08 | ns | | |
| | | | | GT | 44.69 | 1, 12 | <0.0001 | *** | | |
| | | Females | 3-Way ANOVA | Time | 0.068 | 2, 24 | 0.9343 | ns | 1d | n.a. |
| | | | | GT | 108.6 | 1, 24 | <0.0001 | *** | | |
| | | | | ELA | 0.31 | 1, 24 | 0.5844 | ns | | |
| | | | | Time x GT | 0.24 | 2, 24 | 0.7876 | ns | | |
| | | | | Time x ELA | 0.11 | 2, 24 | 0.8959 | ns | | |
| | | | | GT x ELA | 0.8 | 1, 24 | 0.3794 | ns | | |
| | | | | Time x GT x ELA | 0.31 | 2, 24 | 0.7339 | ns | | |
| | | Males | 3-Way ANOVA | Time | 2.17 | 2, 24 | 0.1362 | ns | 1e | n.a. |
| | | | | GT | 92.02 | 1, 24 | <0.0001 | *** | | |
| | | | | ELA | 0.53 | 1, 24 | 0.4751 | ns | | |
| | | | | Time x GT | 0.24 | 2, 24 | 0.7869 | ns | | |
| | | | | Time x ELA | 3.57 | 2, 24 | 0.0441 | * | | |
| | | | | GT x ELA | 0.31 | 1, 24 | 0.5847 | ns | | |
| | | | | Time x GT x ELA | 0.57 | 2, 24 | 0.5749 | ns | | |
| | cFos | P35 Females | 2-Way ANOVA | Interaction | 0.003 | 1, 12 | 0.960 | ns | 1g | n.a. |
| | | | | GT | 2.77 | 1, 12 | 0.120 | ns | | |
| | | | | ELA | 2.58 | 1, 12 | 0.130 | ns | | |
| | | P35 Males | 2-Way ANOVA | Interaction | 0.12 | 1, 12 | 0.739 | ns | 1h | n.a. |
| | | | | GT | 0.43 | 1, 12 | 0.526 | ns | | |
| | | | | ELA | 0.005 | 1, 12 | 0.947 | ns | | |
| | | P56 Females | 2-Way ANOVA | Interaction | 5.164 | 1, 19 | 0.035 | * | 1i | Tukey |
| | | | | GT | 13.37 | 1, 19 | 0.002 | ** | | |
| | | | | ELA | 5.876 | 1, 19 | 0.026 | * | | |
| | | P56 Males | 2-Way ANOVA | Interaction | 0.034 | 1, 17 | 0.855 | ns | 1j | n.a. |
| | | | | GT | 10.12 | 1, 17 | 0.006 | ** | | |
| | | | | ELA | 3.62 | 1, 17 | 0.074 | ns | | |
| | Forced Swim test immobility | P35 Females | 3-Way RM ANOVA | Time | 90.02 | 3.831, 203.1 | <0.0001 | *** | 2b | n.a. |
| | | | | GT | 2.56 | 1, 53 | 0.12 | ns | | |
| | | | | ELA | 0.0004 | 1, 53 | 0.98 | ns | | |
| | | | | Time x GT | 1.09 | 5, 265 | 0.37 | ns | | |
| | | | | Time x ELA | 0.45 | 5, 265 | 0.81 | ns | | |
| | | | | GT x ELA | 0.07 | 1, 53 | 0.80 | ns | | |
| | | | | Time x GT x ELA | 1.19 | 5, 265 | 0.32 | ns | | |
| | | P35 Females average immobility | 2-Way ANOVA | Interaction | 0.017 | 1, 53 | 0.9 | ns | 2c | n.a. |
| | | | | ELA | 0.535 | 1, 53 | 0.47 | ns | | |
| | | | | GT | 3.16 | 1, 53 | 0.08 | ns | | |
| | | | | Litter (covariate) | 1.05 | 1, 53 | 0.31 | ns | | |
| | | P35 Males | 3-Way RM ANOVA | Time | 67.09 | 3.170, 161.7 | <0.0001 | *** | 2e | n.a. |
| | | | | GT | 4.9 | 1, 51 | 0.03 | * | | |
| | | | | ELA | 0.1 | 1, 51 | 0.75 | ns | | |
| | | | | Time x GT | 1.1 | 5, 255 | 0.37 | ns | | |
| | | | | Time x ELA | 0.73 | 5, 255 | 0.60 | ns | | |
| | | | | GT x ELA | 0.15 | 1, 51 | 0.70 | ns | | |
| | | | | Time x GT x ELA | 0.98 | 5, 255 | 0.43 | ns | | |
| | | P35 Males average immobility | 2-Way ANOVA | Interaction | 0.15 | 1, 51 | 0.7 | ns | 2f | n.a. |
| | | | | ELA | 0.15 | 1, 51 | 0.7 | ns | | |
| | | | | GT | 4.85 | 1, 51 | 0.03 | * | | |
| | | | | Litter (covariate) | 0.05 | 1, 51 | 0.83 | ns | | |
| | | P56 Females | 3-Way RM ANOVA | Time | 25.55 | 3.5, 237.7 | <0.0001 | *** | 2i | Tukey |
| | | | | GT | 1.178 | 1, 68 | 0.28 | ns | | |
| | | | | ELA | 0.24 | 1, 68 | 0.62 | ns | | |
| | | | | Time x GT | 1.89 | 5, 340 | 0.10 | ns | | |
| | | | | Time x ELA | 1.24 | 5, 340 | 0.29 | ns | | |

| | | | | | | | | | | |
|---|---|---|---|---|---|---|---|---|---|---|
| | | | | GT x ELA | 10.24 | 1, 68 | 0.002 | ** | | |
| | | | | Time x GT x ELA | 3.42 | 5, 340 | 0.005 | ** | | |
| | | P56 Females average immobility | 2-Way ANOVA | Interaction | 12.93 | 1, 68 | <0.001 | *** | 2j | Tukey |
| | | | | ELA | 1.3 | 1, 68 | 0.260 | ns | | |
| | | | | GT | 0.002 | 1, 68 | 0.970 | ns | | |
| | | | | Litter (covariate) | 2.52 | 1, 68 | 0.12 | ns | | |
| | | P56 Males | 3-Way RM ANOVA | Time | 29.47 | 2.819, 163.5 | <0.0001 | *** | 2l | n.a. |
| | | | | GT | 2.07 | 1, 58 | 0.16 | ns | | |
| | | | | ELA | 0.07 | 1, 58 | 0.79 | ns | | |
| | | | | Time x GT | 0.75 | 5, 290 | 0.59 | ns | | |
| | | | | Time x ELA | 0.49 | 5, 290 | 0.78 | ns | | |
| | | | | GT x ELA | 0.26 | 1, 58 | 0.61 | ns | | |
| | | | | Time x GT x ELA | 0.17 | 5, 290 | 0.97 | ns | | |
| | | P56 Males average immobility | 2-Way ANOVA | Interaction | 0.26 | 1, 58 | 0.62 | ns | 2m | n.a. |
| | | | | ELA | 0.02 | 1, 58 | 0.90 | ns | | |
| | | | | GT | 1.96 | 1, 58 | 0.18 | ns | | |
| | | | | Litter (covariate) | 0.34 | 1, 58 | 0.56 | ns | | |
| | CORT levels | P35 Females | 3-Way RM ANOVA | FST | 18.29 | 1, 36 | 0.0001 | *** | 2d | n.a. |
| | | | | GT | 9.56 | 1, 36 | 0.004 | ** | | |
| | | | | ELA | 0.71 | 1, 36 | 0.40 | ns | | |
| | | | | FST x GT | 1.67 | 1, 36 | 0.20 | ns | | |
| | | | | FST x ELA | 0.7 | 1, 36 | 0.41 | ns | | |
| | | | | GT x ELA | 5.65 | 1, 36 | 0.02 | * | | |
| | | | | FST x GT x ELA | 0.79 | 1, 36 | 0.38 | ns | | |
| | | P35 Males | 3-Way RM ANOVA | FST | 24.44 | 1, 35 | <0.0001 | *** | 2g | n.a. |
| | | | | GT | 0.29 | 1, 35 | 0.59 | ns | | |
| | | | | ELA | 0.02 | 1, 35 | 0.90 | ns | | |
| | | | | FST x GT | 0.02 | 1, 35 | 0.90 | ns | | |
| | | | | FST x ELA | 0.01 | 1, 35 | 0.93 | ns | | |
| | | | | GT x ELA | 0.53 | 1, 35 | 0.47 | ns | | |
| | | | | FST x GT x ELA | 1.07 | 1, 35 | 0.31 | ns | | |
| | | P56 Females | 3-Way RM ANOVA | FST | 73.18 | 1, 36 | <0.0001 | *** | 2k | Tukey |
| | | | | GT | 0.01 | 1, 36 | 0.91 | ns | | |
| | | | | ELA | 2.54 | 1, 36 | 0.12 | ns | | |
| | | | | FST x GT | 0.07 | 1, 36 | 0.79 | ns | | |
| | | | | FST x ELA | 5.77 | 1, 36 | 0.02 | * | | |
| | | | | GT x ELA | 6.08 | 1, 36 | 0.19 | * | | |
| | | | | FST x GT x ELA | 0.87 | 1, 36 | 0.36 | ns | | |
| | | P56 Males | 3-Way RM ANOVA | FST | 77.87 | 1, 38 | <0.0001 | *** | 2n | n.a. |
| | | | | GT | 0.1 | 1, 38 | 0.75 | ns | | |
| | | | | ELA | 0.66 | 1, 38 | 0.42 | ns | | |
| | | | | FST x GT | 0.04 | 1, 38 | 0.84 | ns | | |
| | | | | FST x ELA | 1.44 | 1, 38 | 0.24 | ns | | |
| | | | | GT x ELA | 1.6 | 1, 38 | 0.21 | ns | | |
| | | | | FST x GT x ELA | 2.55 | 1, 38 | 0.12 | ns | | |
| cFos in Paraventricular Nucleus | PVN | P56 Female | 2-Way ANOVA | GT x ELA | 10.89 | 1, 18 | 0.004 | ** | 3c | Tukey |
| | | | | GT | 0.51 | 1, 18 | 0.48 | ns | | |
| | | | | ELA | 0.83 | 1, 18 | 0.37 | ns | | |
| | | P56 Male | 2-Way ANOVA | GT x ELA | 0.14 | 1, 17 | 0.71 | ns | 3d | n.a. |
| | | | | GT | 0.24 | 1, 17 | 0.63 | ns | | |
| | | | | ELA | 2.19 | 1, 17 | 0.16 | ns | | |
| Corticotropin Releasing Factor | CRF/GAPDH | P56 Female | 2-Way ANOVA | GT x ELA | 4.46 | 1, 21 | 0.047 | * | 3e | Tukey |
| | | | | GT | 4.6 | 1, 21 | 0.044 | * | | |
| | | | | ELA | 4.9 | 1, 21 | 0.038 | * | | |
| | | P56 Male | 2-Way ANOVA | GT x ELA | 0.66 | 1, 17 | 0.43 | ns | 3f | n.a. |
| | | | | GT | 0.02 | 1, 17 | 0.90 | ns | | |
| | | | | ELA | 0.26 | 1, 17 | 0.62 | ns | | |
| | | P35 Female vDG | 2-Way ANOVA | GT x ELA | 1.08 | 1, 16 | 0.31 | ns | | n.a. |
| | | | | GT | 1.15 | 1, 16 | 0.30 | ns | | |
| | | | | ELA | 0.77 | 1, 16 | 0.39 | ns | 4c | |
| | | P35 Male vDG | 2-Way ANOVA | GT x ELA | 1.63 | 1,17 | 0.22 | ns | | n.a. |
| | | | | GT | 2.38 | 1,17 | 0.14 | ns | | |
| | | | | ELA | 0.27 | 1,17 | 0.61 | ns | 4d | |
| | | P56 Female vDG | 2-Way ANOVA | GT x ELA | 7.01 | 1, 18 | 0.02 | * | | Tukey |
| | | | | GT | 1.88 | 1, 18 | 0.19 | ns | | |
| | | | | ELA | 13.83 | 1, 18 | 0.002 | ** | 4e | |
| | | P56 Male vDG | 2-Way ANOVA | GT x ELA | 0.25 | 1, 13 | 0.63 | ns | | n.a. |
| | | | | GT | 1.06 | 1, 13 | 0.32 | ns | | |

| | | | | | | | | | | |
|---|---|---|---|---|---|---|---|---|---|---|
| | | | ANOVA | ELA | 0.005 | 1, 13 | 0.94 | ns | 4f | |
| | | P35 Female dDG | 2-Way ANOVA | GT x ELA | 0.13 | 1, 16 | 0.73 | ns | S5c | n.a. |
| | | | | GT | 0.9 | 1, 16 | 0.36 | ns | | |
| | | | | ELA | 3.03 | 1, 16 | 0.10 | ns | | |
| | | P35 Male dDG | 2-Way ANOVA | GT x ELA | 1.35 | 1, 17 | 0.26 | ns | S5d | n.a. |
| | | | | GT | 1.5 | 1, 17 | 0.24 | ns | | |
| | | | | ELA | 3.5 | 1, 17 | 0.08 | ns | | |
| | | P56 Female dDG | 2-Way ANOVA | GT x ELA | 4.26 | 1, 18 | 0.05 | ns | S5e | n.a. |
| | | | | GT | 0.2 | 1, 18 | 0.66 | ns | | |
| | | | | ELA | 3.84 | 1, 18 | 0.07 | ns | | |
| | | P56 Male dDG | 2-Way ANOVA | GT x ELA | 2.01 | 1, 13 | 0.18 | ns | S5f | n.a. |
| | | | | GT | 0.09 | 1, 13 | 0.77 | ns | | |
| | | | | ELA | 0.08 | 1, 13 | 0.78 | ns | | |
| Double Cortin Staining | DCX | P35 vDG | 3-Way ANOVA | GT | 0.07 | 1, 33 | 0.80 | ns | 4c,d | n.a. |
| | | | | Sex | 3.51 | 1, 33 | 0.07 | ns | | |
| | | | | ELA | 0.99 | 1, 33 | 0.33 | ns | | |
| | | | | GT x Sex | 3.38 | 1, 33 | 0.08 | ns | | |
| | | | | GT x ELA | 0.01 | 1, 33 | 0.92 | ns | | |
| | | | | Sex x ELA | 0.09 | 1, 33 | 0.77 | ns | | |
| | | | | GT x Sex x ELA | 2.67 | 1, 33 | 0.11 | ns | | |
| | | P56 vDG | 3-Way ANOVA | GT | 2.8 | 1, 31 | 0.10 | ns | 4e,f | n.a. |
| | | | | Sex | 0.05 | 1, 31 | 0.82 | ns | | |
| | | | | ELA | 3.76 | 1, 31 | 0.06 | ns | | |
| | | | | GT x Sex | 0.04 | 1, 31 | 0.84 | ns | | |
| | | | | GT x ELA | 3.52 | 1, 31 | 0.07 | ns | | |
| | | | | Sex x ELA | 4.27 | 1, 31 | 0.05 | ns | | |
| | | | | GT x Sex x ELA | 0.95 | 1, 31 | 0.34 | ns | | |
| | | P35 dDG | 3-Way ANOVA | GT | 0.02 | 1, 33 | 0.90 | ns | S2c,d | n.a. |
| | | | | Sex | 25.17 | 1, 33 | <0.0001 | *** | | |
| | | | | ELA | 6.53 | 1, 33 | 0.02 | * | | |
| | | | | GT x Sex | 2.33 | 1, 33 | 0.14 | ns | | |
| | | | | GT x ELA | 0.27 | 1, 33 | 0.61 | ns | | |
| | | | | Sex x ELA | 0.0003 | 1, 33 | 0.99 | ns | | |
| | | | | GT x Sex x ELA | 1.1 | 1, 33 | 0.30 | ns | | |
| | | P56 dDG | 3-Way ANOVA | GT | 0.26 | 1, 31 | 0.61 | ns | S2e,f | n.a. |
| | | | | Sex | 0.05 | 1, 31 | 0.82 | ns | | |
| | | | | ELA | 2.51 | 1, 31 | 0.12 | ns | | |
| | | | | GT x Sex | 0.01 | 1, 31 | 0.90 | ns | | |
| | | | | GT x ELA | 5.8 | 1, 31 | 0.02 | * | | |
| | | | | Sex x ELA | 1.44 | 1, 31 | 0.24 | ns | | |
| | | | | GT x Sex x ELA | 0.28 | 1, 31 | 0.60 | ns | | |
| Active Microglia | CD68/Iba1 | P35 Female PVN | 2-Way ANOVA | GT x ELA | 0.48 | 1, 13 | 0.50 | ns | 5c | n.a. |
| | | | | GT | 0.1 | 1, 13 | 0.76 | ns | | |
| | | | | ELA | 0.0006 | 1, 13 | 0.98 | ns | | |
| | | P35 Male PVN | 2-Way ANOVA | GT x ELA | 0.034 | 1, 13 | 0.86 | ns | 5d | n.a. |
| | | | | GT | 0.174 | 1, 13 | 0.68 | ns | | |
| | | | | ELA | 0.561 | 1, 13 | 0.47 | ns | | |
| | | P56 Female PVN | 2-Way ANOVA | GT x ELA | 12.09 | 1, 17 | 0.003 | ** | 5e | Tukey |
| | | | | GT | 0.4 | 1, 17 | 0.53 | ns | | |
| | | | | ELA | 2.4 | 1, 17 | 0.14 | ns | | |
| | | P56 Male PVN | 2-Way ANOVA | GT x ELA | 0.95 | 1, 15 | 0.35 | ns | 5f | n.a. |
| | | | | GT | 1.8 | 1, 15 | 0.20 | ns | | |
| | | | | ELA | 1.76 | 1, 15 | 0.21 | ns | | |
| | | P35 Female vDG | 2-Way ANOVA | GT x ELA | 2.79 | 1, 20 | 0.11 | ns | 5g | n.a. |
| | | | | GT | 12.21 | 1, 20 | 0.002 | ** | | |
| | | | | ELA | 0.5 | 1, 20 | 0.49 | ns | | |
| | | P35 Male vDG | 2-Way ANOVA | GT x ELA | 3.15 | 1, 17 | 0.09 | ns | 5h | n.a. |
| | | | | GT | 1.32 | 1, 17 | 0.27 | ns | | |
| | | | | ELA | 0.02 | 1, 17 | 0.88 | ns | | |
| | | P56 Female vDG | 2-Way ANOVA | GT x ELA | 11.74 | 1, 21 | 0.003 | ** | 5i | Tukey |
| | | | | GT | 5.91 | 1, 21 | 0.024 | * | | |
| | | | | ELA | 0.15 | 1, 21 | 0.70 | ns | | |
| | | P56 Male vDG | 2-Way ANOVA | GT x ELA | 2.6 | 1, 17 | 0.13 | ns | 5j | n.a. |
| | | | | GT | 0.79 | 1, 17 | 0.39 | ns | | |
| | | | | ELA | 0.65 | 1, 17 | 0.43 | ns | | |
| | | P35 Female dDG | 2-Way ANOVA | GT x ELA | 0.24 | 1, 20 | 0.63 | ns | S6c | n.a. |
| | | | | GT | 7.7 | 1, 20 | 0.01 | * | | |
| | | | | ELA | 4.39 | 1, 20 | 0.049 | * | | |

| | | | | | | | | | | |
|---|---|---|---|---|---|---|---|---|---|---|
| | | P35 Male dDG | 2-Way ANOVA | GT x ELA | 0.18 | 1, 17 | 0.67 | ns | | n.a. |
| | | | | GT | 3.2 | 1, 17 | 0.09 | ns | | |
| | | | | ELA | 1.47 | 1, 17 | 0.24 | ns | S6d | |
| | | P56 Female dDG | 2-Way ANOVA | GT x ELA | 2.49 | 1, 20 | 0.13 | ns | | n.a. |
| | | | | GT | 0.09 | 1, 20 | 0.77 | ns | | |
| | | | | ELA | 0.51 | 1, 20 | 0.48 | ns | S6e | |
| | | P56 Male dDG | 2-Way ANOVA | GT x ELA | 1.03 | 1, 18 | 0.32 | ns | | n.a. |
| | | | | GT | 0.03 | 1, 18 | 0.87 | ns | | |
| | | | | ELA | 0.022 | 1, 18 | 0.89 | ns | S6f | |
| Total microglia | Iba1 | P35 Female PVN | 2-Way ANOVA | GT x ELA | 0.3 | 1, 13 | 0.59 | ns | | n.a. |
| | | | | GT | 0.04 | 1, 13 | 0.84 | ns | | |
| | | | | ELA | 0.51 | 1, 13 | 0.49 | ns | S7a | |
| | | P35 Male PVN | 2-Way ANOVA | GT x ELA | 4.01 | 1, 13 | 0.07 | ns | | n.a. |
| | | | | GT | 0.03 | 1, 13 | 0.87 | ns | | |
| | | | | ELA | 2.37 | 1, 13 | 0.09 | ns | S7b | |
| | | P56 Female PVN | 2-Way ANOVA | GT x ELA | 0.11 | 1, 17 | 0.74 | ns | | n.a. |
| | | | | GT | 0.17 | 1, 17 | 0.07 | ns | | |
| | | | | ELA | 1.6 | 1, 17 | 0.22 | ns | S7c | |
| | | P56 Male PVN | 2-Way ANOVA | GT x ELA | 3.61 | 1, 15 | 0.08 | ns | | Tukey |
| | | | | GT | 3.51 | 1, 15 | 0.08 | ns | | |
| | | | | ELA | 4.84 | 1, 15 | 0.04 | * | S7d | |
| | | P35 Female vDG | 2-Way ANOVA | GT x ELA | 0.003 | 1, 20 | 0.96 | ns | | n.a. |
| | | | | GT | 2.77 | 1, 20 | 0.11 | ns | | |
| | | | | ELA | 0.0002 | 1, 20 | 0.99 | ns | S7e | |
| | | P35 Male vDG | 2-Way ANOVA | GT x ELA | 3.95 | 1, 17 | 0.06 | ns | | n.a. |
| | | | | GT | 0.00004 | 1, 17 | 1.00 | ns | | |
| | | | | ELA | 5.96 | 1, 17 | 0.03 | * | S7f | |
| | | P56 Female vDG | 2-Way ANOVA | GT x ELA | 0.05 | 1, 19 | 0.82 | ns | | n.a. |
| | | | | GT | 0.03 | 1, 19 | 0.86 | ns | | |
| | | | | ELA | 3.31 | 1, 19 | 0.08 | ns | S7g | |
| | | P56 Male vDG | 2-Way ANOVA | GT x ELA | 0.0007 | 1, 21 | 0.98 | ns | | n.a. |
| | | | | GT | 3.13 | 1, 21 | 0.09 | ns | | |
| | | | | ELA | 0.33 | 1, 21 | 0.57 | ns | S7h | |
| | | P35 Female dDG | 2-Way ANOVA | GT x ELA | 0.01 | 1, 20 | 0.91 | ns | | n.a. |
| | | | | GT | 0.54 | 1, 20 | 0.47 | ns | | |
| | | | | ELA | 0.05 | 1, 20 | 0.82 | ns | S7i | |
| | | P35 Male dDG | 2-Way ANOVA | GT x ELA | 6.03 | 1, 16 | 0.03 | * | | Tukey |
| | | | | GT | 6.24 | 1, 16 | 0.02 | * | | |
| | | | | ELA | 6.71 | 1, 16 | 0.02 | * | S7j | |
| | | P56 Female dDG | 2-Way ANOVA | GT x ELA | 0.07 | 1, 20 | 0.79 | ns | | n.a. |
| | | | | GT | 1.2 | 1, 20 | 0.29 | ns | | |
| | | | | ELA | 0.0004 | 1, 20 | 0.98 | ns | S7k | |
| | | P56 Male dDG | 2-Way ANOVA | GT x ELA | 0.91 | 1, 18 | 0.35 | ns | | n.a. |
| | | | | GT | 0.17 | 1, 18 | 0.69 | ns | | |
| | | | | ELA | 0.91 | 1, 18 | 0.35 | ns | S7l | |
| 5HT1A KD | 5HT1A/TPH2 | Male vs Female | 3-Way RM ANOVA | Time | 0.63 | 2.299, 18.40 | 0.5669 | ns | S1 | n.a. |
| | | | | GT | 147.3 | 1, 8 | <0.0001 | *** | | |
| | | | | Sex | 0.69 | 1, 8 | 0.4318 | ns | | |
| | | | | Time x GT | 1.7 | 3, 24 | 0.1946 | ns | | |
| | | | | Time x Sex | 0.63 | 3, 24 | 0.6036 | ns | | |
| | | | | GT x Sex | 5.24 | 1, 8 | 0.0513 | ns | | |
| | | | | Time x GT x Sex | 1.37 | 3, 24 | 0.2748 | ns | | |
| Baseline CORT P35 vs P56 | CORT | Females | 2-Way ANOVA | Age x ELA | 3.52 | 1, 44 | 0.067 | ns | S2a | Tukey |
| | | | | Age | 15.72 | 1, 44 | 0.0003 | *** | | |
| | | | | ELA | 12.1 | 1, 44 | 0.001 | ** | | |
| | | Males | 2-Way ANOVA | Age x ELA | 0.055 | 1, 40 | 0.82 | ns | S2b | Tukey |
| | | | | Age | 8.75 | 1, 40 | 0.005 | ** | | |
| | | | | ELA | 0.007 | 1, 40 | 0.93 | ns | | |
| Body weight | g | Female | -Way ANOV/ | Age | 365.5 | 3, 195 | <0.0001 | **** | S3b | n.a. |
| | | | | Gentoype | 0.36 | 1, 195 | 0.55 | ns | | |
| | | | | ELA | 8.88 | 1, 195 | 0.00 | ** | | |
| | | | | Age x Gentoype | 0.1 | 3, 195 | 0.96 | ns | | |
| | | | | Age x ELA | 9.63 | 3, 195 | <0.0001 | **** | | |
| | | | | Gentoype x ELA | 0.00002284 | 1, 195 | 1.00 | ns | | |
| | | | | Age x Gentoype x I | 0.13 | 3, 195 | 0.94 | ns | | |
| | | Male | -Way ANOV/ | Age | 503.7 | 3, 195 | <0.0001 | **** | S3c | n.a. |
| | | | | Gentoype | 1.27 | 1, 195 | 0.26 | ns | | |
| | | | | ELA | 16.94 | 1, 195 | <0.0001 | **** | | |
| | | | | Age x Gentoype | 0.12 | 3, 195 | 0.95 | ns | | |

| | | | | | | | | | | |
|---|---|---|---|---|---|---|---|---|---|---|
| | | | | Age x ELA | 3.21 | 3, 195 | 0.02 | * | | |
| | | | | Gentoype x ELA | 2.42 | 1, 195 | 0.12 | ns | | |
| | | | | Age x Gentoype x I | 0.87 | 3, 195 | 0.46 | ns | | |
| Avoidance Behavior | EPM Open Arm Time | P56 Female | 2-Way ANOVA | GT x ELA | 0.68 | 1, 32 | 0.415 | ns | S4b | n.a |
| | | | | GT | 1.464 | 1, 32 | 0.235 | ns | | |
| | | | | ELA | 0.737 | 1, 32 | 0.396 | ns | | |
| | | P56 Male | 2-Way ANOVA | GT x ELA | 1.31 | 1, 35 | 0.26 | ns | S4c | n.a |
| | | | | GT | 0.016 | 1, 35 | 0.9 | ns | | |
| | | | | ELA | 3.59 | 1, 35 | 0.07 | ns | | |
